# Cold seeps are hotspots of deep-sea nitrogen-loss driven by microorganisms across 21 phyla

**DOI:** 10.1101/2024.06.05.597523

**Authors:** Qiuyun Jiang, Lei Cao, Yingchun Han, Shengjie Li, Rui Zhao, Xiaoli Zhang, S. Emil Ruff, Zhuoming Zhao, Jiaxue Peng, Jing Liao, Baoli Zhu, Minxiao Wang, Xianbiao Lin, Xiyang Dong

## Abstract

Nitrogen bioavailability, governed by the balance of fixation and loss processes, is a key factor regulating oceanic productivity, ecosystem functions, and global biogeochemical cycles. The key nitrogen-loss organisms—denitrifiers and anaerobic ammonium-oxidizing (anammox) bacteria—are not well understood in marine seafloor environments, especially in deep-sea cold seeps. In this study, we combined geochemical measurements, ^15^N stable isotope tracer analysis, metagenomics, metatranscriptomics, and three-dimensional protein structural simulations to investigate the diversity of denitrifying and anammox microbial communities and their biogeochemical roles in these habitats. Geochemical evidence from 301 sediment samples shows significantly higher nitrogen-loss rates in cold seeps compared to typical deep-sea sediments, with an estimated annual nitrogen loss of 6.16 Tg from seafloor surface sediments. Examination of a total of 147 million non-redundant genes reveals a high prevalence and active expression of nitrogen-loss genes, including nitrous-oxide reductase (NosZ; 6.88 genes per million or GPM on average), nitric oxide dismutase (Nod; 1.29 GPM), and hydrazine synthase (HzsA; 3.35 GPM) in surface sediments. Analysis of 3,164 metagenome-assembled genomes from this habitat has expanded the known diversity of nitrous-oxide reducers to six phyla and nitric oxide-dismutating organisms to one phylum and two new orders, while ten phyla host anammox bacteria going beyond *Planctomycetota*. These microbes show diverse structural adaptations and complex gene cluster arrangements that potentially enable survival in the harsh conditions of cold seeps. These findings suggest that cold seeps, despite their low temperatures, are significant, previously underestimated hotspots of nitrogen loss, potentially contribute substantially to the global nitrogen cycle.

## Introduction

Cold seeps are specialized marine environments primarily located along continental slopes and subduction zones, where subsurface fluids rich in hydrogen sulfide and hydrocarbons, such as methane, seep through the seabed^1–3^. These environments support complex ecosystems centered on the anaerobic oxidation of methane (AOM), a process involving methane-consuming archaea and sulfate-reducing bacteria. All life in these deep-sea oases depends on bioavailable nitrogen to support growth, which is a critical factor limiting biological productivity. Thus, understanding the processes that balance the nitrogen budget in cold seeps is essential^4, 5^. Diazotrophs, organisms that convert dinitrogen gas (N_2_) into bioavailable nitrogen, including e.g. ammonium, nitrite, and nitrate, through biological nitrogen fixation, are widespread in these environments, supported by diverse energy sources from either cultivated or uncultivated lineages^6^. Concurrently, nitrogen-loss microbes convert bioavailable nitrogen back into N_2_ to maintain a balanced nitrogen cycle. However, studies on nitrogen-loss processes in deep-sea cold seeps and the responsible microbial communities are relatively limited^7–9^. Understanding these processes is crucial for comprehending how cold seeps contribute to the broader nitrogen cycle in marine systems.

In marine sediments, two primary microbial processes for nitrogen-loss are denitrification and anaerobic ammonium oxidation (anammox)^10, 11^. Denitrification occurs in two forms: classical and oxygenic. Classical denitrification reduces nitrate (NO_3_^−^) to nitrite (NO_2_^−^) and then to nitric oxide (NO), which is subsequently converted to nitrous oxide (N_2_O). This N_2_O is further reduced to N_2_ by the enzyme nitrous-oxide reductase (N_2_OR), encoded by the *nosZ* gene cluster^12^. Oxygenic denitrification simplifies this process by converting NO into N_2_ and molecular oxygen (O_2_) through the action of nitric oxide dismutase (Nod), bypassing the production of N_2_O^13, 14^. Anammox, on the other hand, combines NO_2_ ^−^ with ammonium (NH_4_ ^+^) to form N_2_, providing a more energy-efficient pathway for nitrogen removal without relying on oxygen^10, 15^. This process includes the reduction of NO_2_^−^ to NO, which then reacts with NH_4_^+^ to create hydrazine (N_2_H_4_), a highly reactive and toxic compound with a low redox potential which then is oxidized to N_2_. The hydrazine-forming reaction, facilitated by hydrazine synthase (Hzs) is essential for the anammox pathway due to its unique biochemical properties.

The relative contributions of denitrification and anammox to nitrogen-loss vary across different marine sediments. Most studies have focused on areas within 1000 meters of water depth, where denitrification typically accounts for over 80% of nitrogen-loss. Denitrification rates in estuarine and coastal environments show a broad range (0– 217.9 nmol cm^-3^ h^-1^)^11, 16–21^. In contrast, rates in continental shelves, slopes, and deep-sea cold seep sediments vary from 0–82.97 mmol m^-3^ h^-1^ ^17, 19, 22–25^. Meanwhile, in deeper sea sediments (e.g., trenches), anammox contributes to about 50% of nitrogen-loss. Studies by Thamdrup et al.^11^ in Atacama and Kermadec Trenches show that anammox bacteria are the main drivers of nitrogen loss in these deep-sea environments, whereas denitrification occurs at slower rates and is confined to the surface layers, which are potentially influenced by water depth and the scarcity of reactive organic carbon^11, 26, 27^. Although anammox plays a dominant role in deep-sea sediments, denitrification can still predominate under certain conditions, particularly in sediments with water depths greater than 1000 meters, where it contributes between 40.73% and 88.06% to nitrogen loss^22, 25, 27^. These observations indicate that both denitrification and anammox may play a role in nitrogen loss at cold seeps. However, the exact contributions of these processes in balancing the nitrogen budget at cold seep sites remain unclear.

N_2_OR, encoded by *nosZ* clade I and II, is the only enzyme that biologically converts N_2_O to N_2_ ^28^. NosZ clade I is primarily found in some members of *Alpha*-, *Beta*-, and *Gammaproteobacteria* that possess the complete denitrification pathway. In contrast, NosZ clade II has been found in diverse bacterial groups that lack nitrite reductase genes (*nirS* or *nirK*), including *Gemmatimonadetes*, *Verrucomicrobia*, *Gammaproteobacteria*, and *Chloroflexi*^29–31^. While NosZ clade I is well-documented, recent studies suggest that NosZ clade II may play a more important role in reducing N_2_O in certain ecosystems than previously assumed^29, 31–33^. The *nod* genes were first identified in the genomes of *Candidatus* Methylomirabilis oxyfera-like bacteria (also known as NC10 or nitrite-dependent methane-oxidizing bacteria)^14, 34, 35^, which use Nod to generate N_2_ and O_2_. The generated O_2_ is utilized to catalyze methane oxidation, thereby coupling oxygenic denitrification with aerobic methanotrophic pathways in anoxic environments^34, 36, 37^. These organisms have been detected in globally distributed cold seeps and other seafloor sediments through metagenomic and amplicon sequencing of *pmoA* or 16S rRNA genes^38–40^. Other bacteria like the *Gammaproteobacterium* strain *HdN1*, and species from the genera *Sediminibacterium* and *Algoriphagus* (within the phylum *Bacteroidota*)^41^, which also possess *nod* genes, may produce oxygen via dismutation, suggesting the potential involvement of other microbes in oxygen production in cold seeps.

Known anammox bacteria are all affiliated with the *Planctomycetota* phylum, specifically within five families in the *Brocadiales* order: *Candidatus* Scalinduaceae, *Candidatus* Brocadiaceae, *Candidatus* Anammoxibacteraceae, *Candidatus* Bathyanammoxibiaceae, and *Candidatus* Subterrananammoxibiaceae^42–44^. Although these lineages are widely distributed across marine ecosystems and found in sediments from diverse marine environments, pure cultures have not yet been obtained^27, 45, 46^. Further studies using amplicon sequencing of 16S rRNA, hydrazine dehydrogenase (*hzo*), and *hzsB* genes have discovered diverse anammox bacteria in deep-sea cold seep sediments of the Okhotsk Sea and the South China Sea^8, 9^. These findings suggest that the deep-sea cold seep environment might harbor novel anammox bacteria outside the *Planctomycetota* phylum.

In this study, we aim to investigate the contributions of denitrifying and anammox microbial communities in nitrogen-loss at cold seep habitats, along with their diversity. We first provide geochemical evidence for nitrogen loss in cold seeps, based on data from 301 sediment samples collected from three sites—Lingshui, Haima, and Site F (**Fig. 1 and Supplementary Fig. 1**). This evidence is further supported by measurements of denitrification and anammox activities, conducted through slurry incubation experiments with ^15^N-labelled tracers. Subsequently, we explore the genes associated with nitrogen loss (*nosZ*, *nod*, and *hzsA*) and the diversity of microbes linked to this process. This is achieved through sequence- and structure-based analyses using a detailed gene and genome catalogue compiled from 165 metagenomes from 16 cold seep sites. Our findings reveal that cold seeps are overlooked areas for nitrogen loss in marine sediments under high pressure and low temperature. Nitrogen-loss in this habitat is mediated by diverse microbial populations, including newly identified phyla of anammox bacteria, contributing to this process.

**Figure 1.**
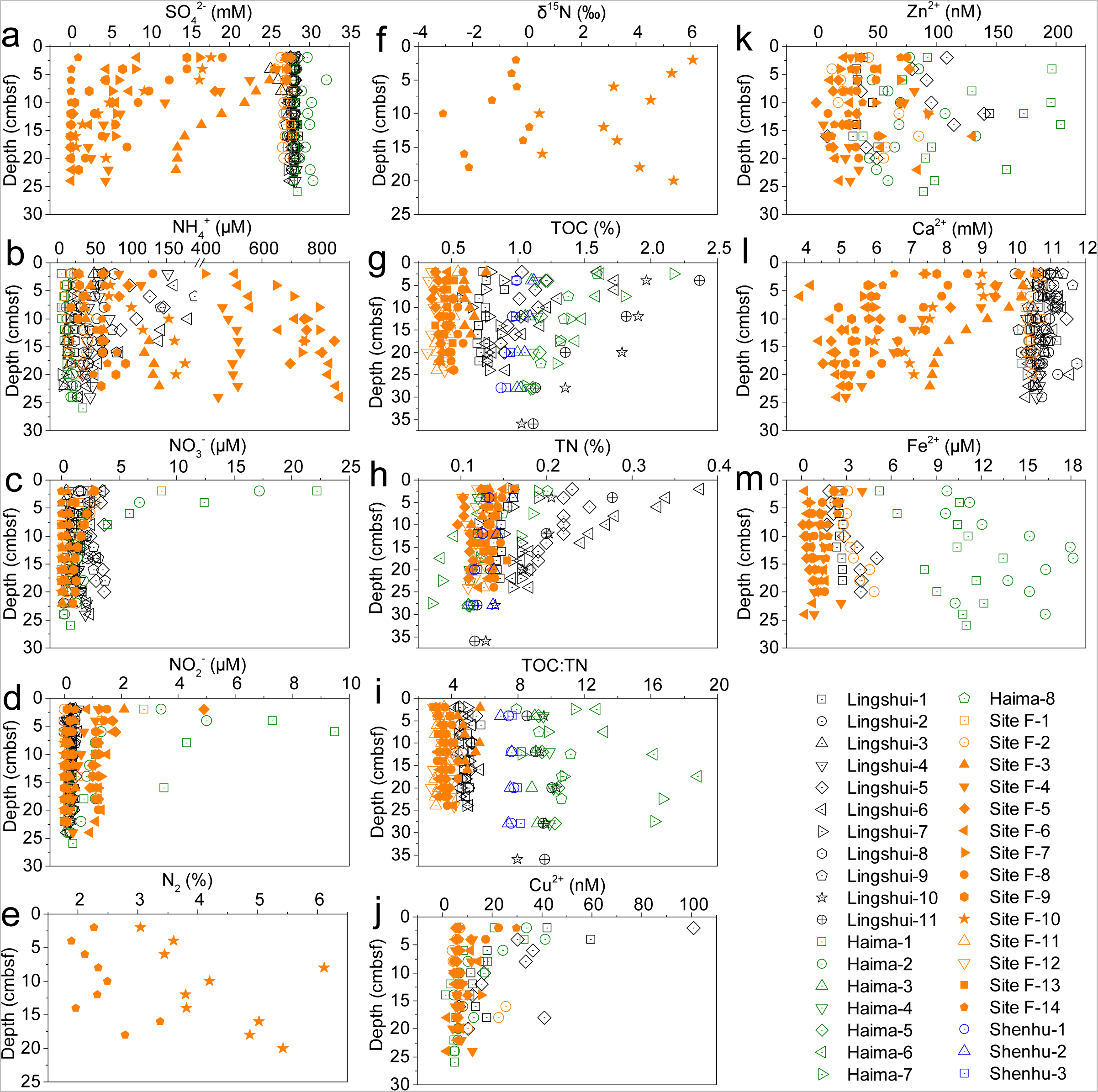
Geochemical characteristics of sediment and porewater in the cold seeps. (a-d) Depth concentration profiles of nutrients (SO_4_^2-^, NH_4_^+^, NO_3_^-^ and NO_2_^-^) in porewater (0-26 cmbsf, n = 221) collected from Lingshui, Haima and Site F cold seeps. (e-f) Depth profiles of N_2_ and their δ^15^N values in the headspace gas of Site F-10 and Site F-14 cold seep sediments. (g-i) Depth profiles of TOC, TN and TOC: TN in cold seep sediments (0-36 cmbsf, n = 163) collected from Lingshui, Haima, and Site F cold seeps. (j-m) Depth concentration profiles of dissolved metals (Cu^2+^, Zn^2+^, Ca^2+^ and Fe^2+^) in porewater (0-26 cmbsf, n = 221) collected from Lingshui, Haima and Site F cold seeps. Black, green, orange and dark blue points represent the concentration of geochemical parameters in Lingshui cold seep, Haima cold seep, Site F cold seep and Shenhu non-seep, respectively. Detailed data can be found in **Supplementary Table 1.**

## Results and discussion

### Geochemical evidence for nitrogen-loss in cold seeps

A total of 301 sediment samples from 33 cores with depths of 0–36 cm below the seafloor (cmbsf) were collected from three different cold seeps—Lingshui, Haima, and Site F (**Supplementary Fig. 1**). These sites represent different stages of cold seep activity with varying methane seeping intensity^47^. Correspondingly, sulfate (SO_4_^2–^) concentrations in the porewater showed coherent downcore variations, either decreasing dramatically or remaining relatively stable depending on methane fluxes (**Fig. 1a and Supplementary Table 1**). The predominant form of dissolved inorganic nitrogen was ammonium (NH_4_^+^; **Fig. 1b**), with the highest concentrations found at Site F (8.6–867.2 μM, average 266.0 μM) and lower concentrations at Lingshui and Haima (averaged at 61.5 and 15.0 μM, respectively). Porewater nitrate (NO_3_^−^) and nitrite (NO_2_^−^) concentrations (averaged at 1.6 and 0.6 μM, respectively) were generally lower than NH_4_^+^ across all sites (**Fig. 1c-d**). Following this, we detected the signature of active N_2_ production, as evident by the downcore increasing concentrations of N_2_ (1.89–6.12%) in two cores, with δ^15^N values in the range of −3.08–6.11‰ (**Fig. 1e-f**). The δ^15^N values of N_2_ indicated that Site F-14 (−3.09–0.10‰) suggest intensive denitrification and anammox activities, indicating significant nitrogen consumption through these processes^48, 49^. Together, our extensive geochemical measurements indicate the occurrence of nitrogen loss in cold seep sediments.

Total organic carbon (TOC), total nitrogen (TN), and TOC:TN ratio varied considerably between sites, generally decreasing with sediment depth (**Fig. 1g-i**). TOC levels in Lingshui, Haima, and Shenhu (non-seep; n = 12) sites were much higher than those in Site F. Enzymes involved in microbial nitrogen-loss processes require metal cofactors^50–52^, which were abundant in all studied sediments. In Lingshui, copper (Cu^2+^) reached the highest concentration, averaging 26.6 nM, while zinc (Zn^2+^) peaked in Haima, averaging 100.4 nM (**Fig. 1j-k**). Calcium (Ca^2+^) displayed a decreasing trend in Site F, while iron (Fe^2+^) was highest in Haima, averaging 11.9 μM (**Fig. 1l-m**).

### Significant potential nitrogen-loss rates in cold seeps

We examined potential nitrogen-loss rates and the contribution of anammox to N_2_ production by conducting slurry experiments with nitrogen isotope tracing on sediment samples up to 36 cm deep from Lingshui, Haima, and Shenhu (**Supplementary Fig. 1**) at 4 °C. We detected both denitrification and anammox, with rates significantly higher in cold seep regions (n = 37) compared to non-seep regions (n = 11; *P* < 0.0001; **Fig. 2b and Supplementary Table 2**). In cold seeps, average potential denitrification rates were measured at 2.81 ± 4.53 nmol cm^-3^ h^-1^, and anammox rates were 0.17 ± 0.25 nmol cm^-3^ h^-1^. In contrast, non-seep regions had denitrification and anammox rates of only 0.1 ± 0.07 nmol cm^-3^ h^-1^ and 0.02 ± 0.02 nmol cm^-3^ h^-1^, respectively. It should be noted that the lesser importance of anammox (18.3 ± 12.73%) in the non-seep sediments is mainly attributed to higher organic carbon contents (0.99 ± 0.08%) (**Fig. 1g**), which favor denitrification over anammox^11, 22^.

**Figure 2.**
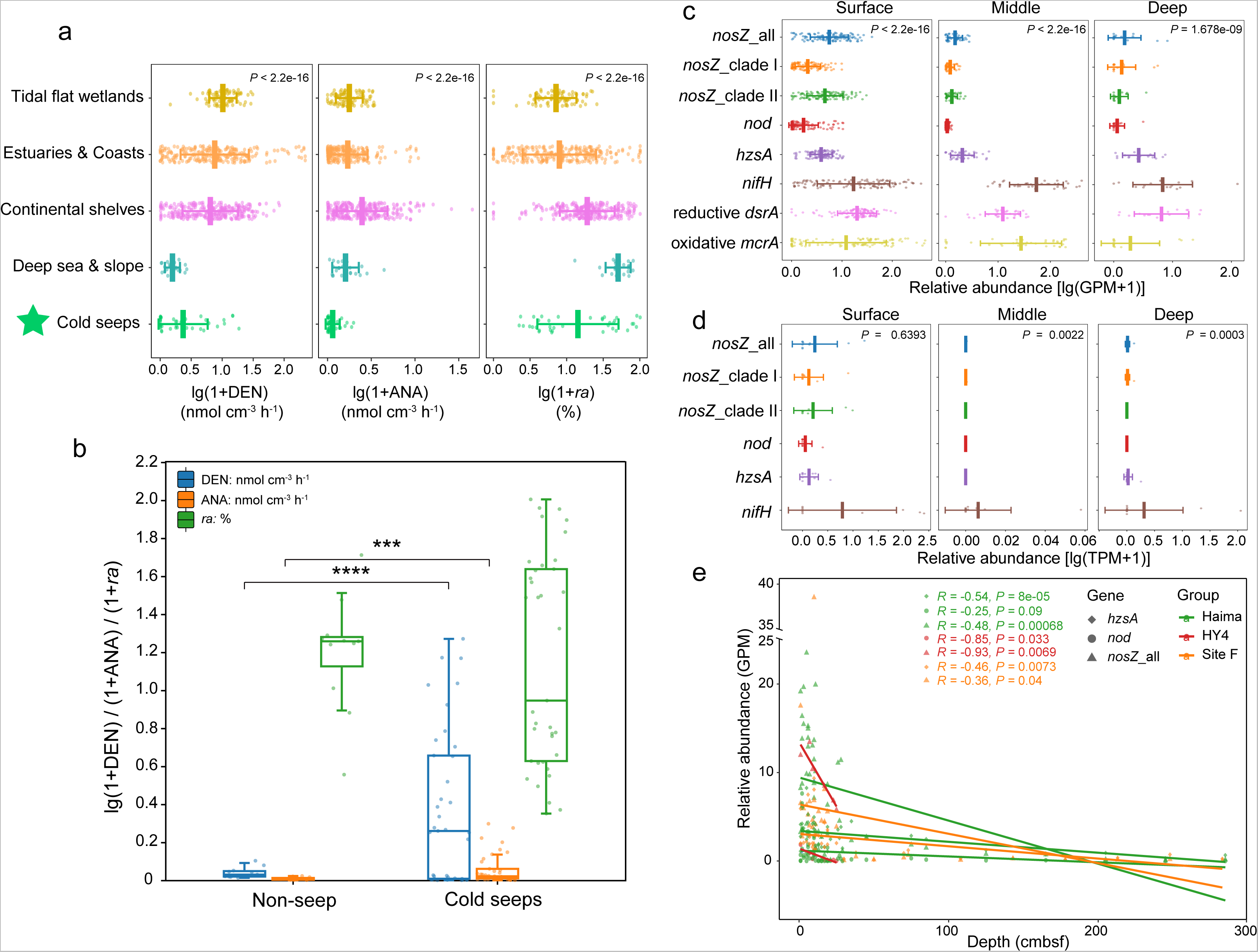
Potential N_2_ production rates in sediments from various environments and relative abundance patterns of nitrogen-loss genes in cold seep sediments. (a) Potential N_2_ production rates (denitrification and anammox) and the percentage of N_2_ production attributed to anammox (*ra*, %) in sediments from various environments. DEN, ANA and *ra* represent denitrification rates, anammox rates, and the proportion of N_2_ produced by anammox, respectively. *P* values for differences across environments were computed using Kruskal-Wallis rank-sum tests. (b) Potential N_2_ production rates (denitrification and anammox) and the percent of N_2_ production attributed to anammox (*ra*, %) in non-seep and cold seeps sediments. *P* values for differences between non-seep and cold seeps were computed using student’s *t*-test. (c) Relative abundance of *nosZ*, *nod*, *hzsA*, *nifH*, reductive *dsrA* and oxidative *mcrA* genes across different sediment samples, measured as genes per million (GPM) for metagenomes. Inserted plots show the abundance of nitrogen-loss genes: all *nosZ* genes (dark blue), *nosZ* Clade I genes (orange), *nosZ* Clade II genes (green), *nod* genes (dark red), *hzsA* genes (dark purple), *nifH* genes (brown), reductive *dsrA* genes (pink), and oxidative *mcrA* genes (dark yellow). *P* values for differences across genes were computed using Kruskal-Wallis rank-sum tests. (d) Expression levels of *nosZ*, *nod*, *hzsA* and *nifH* genes across different sediment samples, measured as transcripts per million (TPM) for metatranscriptomes. Inserted plots show the transcriptional abundance of nitrogen-loss genes. *P* values for differences across genes were computed using Kruskal-Wallis rank-sum tests. (e) Relationships between relative abundance (GPM) of genes (*nosZ*, *nod*, *hzsA*) and depths (cmbsf) of cold seep sediment samples. Each point represents the average gene abundance for a sample, with linear regression lines and *R* values for each gene group shown in corresponding colors. Detailed data are provided in **Supplementary Tables 2-5**.

Although the anammox contributions measured here were lower than that in many previous studies in deep-sea sediments (**Fig. 2a**), they were comparable to values measured in sediments from continental slope^25, 27^ and deep-sea^26^ environments. Denitrification is the primary nitrogen-loss process in these environments, with anammox contributing to roughly 27.44 ± 29.96% of the total nitrogen-loss in cold seeps (**Supplementary Fig. 2**). The rich organic carbon and tight correlation between denitrification rates and TOC contents in cold seeps (*P* < 0.01; **Supplementary Fig. 3a**) suggest that heterotrophic denitrification is a likely metabolic pathway^7^. A tight correlation between anammox and denitrification rates was both observed in cold seeps and non-seep (*P* < 0.01; **Supplementary Fig. 3c and d**), aligning with the fact that denitrification from nitrate to nitrite provides the necessary NO_2_^−^ for anammox in marine sediments^53, 54^. Despite low-temperature conditions, several cold seep sites— specifically Haima-6, Haima-7, Haima-8, Lingshui-10, and Lingshui-11—exhibited exceptionally high rates of nitrogen loss (up to 17.65 ± 1.24 nmol cm^-3^ h^-1^; **Supplementary Fig. 2 and Supplementary Table 2**) possibly related to abundant carbon sources (1.01–2.37%; **Supplementary Table 1**). These rates are comparable to those in estuarine and coastal environments with higher temperatures, as well as other known cold seep sites such as those in the Gulf of Mexico, but are considerably higher than typical deep-sea sediments (**Figure 2a and Supplementary Table 3**).

These findings suggest that certain denitrifying or anammox microbes have adapted or evolved to thrive in deep-sea cold seeps with low temperature, high hydrostatic pressure, and stable carbon and energy supply. Although our experiments did not replicate the high hydrostatic pressure found in natural settings, research has shown that N_2_ production rates measured in the laboratory closely matched those observed *in situ* using a benthic lander in hadal sediments of the Atacama Trench^11^. We extrapolated the potential nitrogen-loss rates measured in our study across the globally estimated seepage area using methods from a previous study^24^. Given that the total active seep areas in Haima is about 350 km^2^ ^55^ and the sediment mean bulk dry density of about 1.3 g cm^-3^ ^56^, and considering the more than 900 global cold seeps^57^, we estimate the global cold seep area to be approximately 3.15×10^5^ km^2^ The estimated nitrogen-loss flux from denitrification and anammox in the surface sediments (0–5 cm) of cold seeps is around 6.16 Tg N per year. This represents about 2.05% of the global marine sediment nitrogen-loss flux (300 Tg N per year)^58^, highlighting that cold seeps, despite covering only about 0.087% of the global marine area, are significant nitrogen-loss hotspots in marine sediments. However, it should be noted that cold seep areas are not yet precisely described in the studies, and this estimate is merely possible, as cold seep areas can vary significantly in size.

### Diverse nitrogen-loss genes are mainly found to be abundant in surface sediments

Using a gene catalog of 147 million non-redundant genes from cold seeps^59^, we delved into the diversity of genes linked to nitrogen loss, focusing on nitrous-oxide reductase (NosZ), nitric oxide dismutase (Nod), and hydrazine synthase (HzsA) (**Supplementary Fig. 4**). We identified 530 NosZ sequences containing cupredoxin-related protein active domains (**Supplementary Fig. 5**). These sequences are categorized into two groups based on their signal peptides: clade I (n = 164) utilizes the twin-arginine translocation (Tat) pathway^60^, while clade II (n = 366) has an additional *c*-type heme domain at the C-terminus, associated with the secretory (Sec) pathway^30, 33^. Furthermore, we discovered a sequence-divergent branch called NosZG7 (n = 27), which, despite sequence differences, showed structural congruence with canonical NosZ (**Supplementary Fig. 5b**). In addition, we identified 151 Nod sequences, all bearing conserved catalytic site residues similar to those of nitric oxide dismutase^41^ from the NO-dismutating bacterium *Methylomirabilis oxyfera* (**Supplementary Fig. 6**). The sequence-divergent branches Cluster1 (n = 28) and Cluster5 (n = 8) were only found through phylogenetic and structural analysis, respectively. We also identified 644 HzsA sequences with hydrazine synthase alpha subunit domains and the pentacoordinated *c*-type heme^52^. The HzsA sequences were classified into six clades, with five branches identified through structural analysis as divergent from those of known anammox bacteria (**Supplementary Fig. 7**). Notably, while no *hzsA* and *nod* protein sequences were detected in mobile genetic elements (MGEs), we found 31 *nosZ* sequences in MGEs, with clade I (n = 24; **Supplementary Fig. 8**) being more prevalent than clade II (n = 7). This suggests that horizontal gene transfer via MGEs may contribute to the diversification of *nosZ*- bearing denitrifying microorganisms in deep-sea cold seeps^61^.

The average abundances of *nosZ*, *nod*, and *hzsA* genes were 4.84, 0.90 and 2.78 genes per million (GPM; 0–6855 cmbsf; **Fig. 2c, Supplementary Fig. 9a and Supplementary Table 4**), respectively. These abundances are one-quarter that of the reductive *dsrA* gene (averaging 21.32 GPM), indicative of sulfate reduction, and only one-tenth of the oxidative *mcrA* gene (averaging 46.15 GPM), which is indicative of methane oxidation. Similarly, they are one-tenth of the *nifH* gene (averaging 55.45 GPM), associated with nitrogen fixation. This imbalance between genes for nitrogen fixation and nitrogen loss suggests that there might be additional pathways for nitrogen loss or the active assimilation of ammonium. The expression levels of nitrogen-loss genes indicate that microbial nitrogen-loss processes are active, particularly at the surface of cold seep sediments, with average values of 2.38 transcripts per million (TPM) for *nosZ*, 0.20 TPM for *nod*, and 0.48 TPM for *hzsA* (**Fig. 2d, Supplementary Fig. 9b and Supplementary Table 5**). Specifically, *nosZ* clade II genes were more abundant than clade I genes (*P* < 0.0001; **Fig. 2c, Supplementary Fig. 9a and Supplementary Table 4**). The expression level of *nosZ* clade II genes (averaging 3.71 TPM) was also higher than that of *nosZ* clade I genes (averaging 1.13 TPM; *P* < 0.0001; **Supplementary Fig. 9b**), indicating that *nosZ* clade II may play a more important role in N_2_O consumption in cold seeps ^29, 32, 33^. The distributions of *nosZ*, *nod*, and *hzsA* genes varied with sediment depth (0–300 cmbsf), showing generally negative trends with increasing depth, indicating diminished nitrogen-loss activity deeper in the sediments (**Fig. 2e**). However, within the shallow surface layers (up to 40 cmbsf), the abundance of *nosZ*, *nod*, and *hzsA* genes positively correlated with depth (*P* < 0.05; **Supplementary Fig. 9c and Supplementary Table 7**). Additionally, cold seep nitrogen-loss gene abundance exhibited statistically significant differences across surface and middle/deep depths, decreasing with sediment depth, with lower abundances in deeper sediments. Specifically, gene abundances ranged from 1.29 to 6.88 GPM at the surface (0–50 cmbsf), 0.08 to 1.41 GPM in the middle depth (50–500 cmbsf), and 0.20 to 0.98 GPM in deep sediments (>500 cmbsf) (*P* < 0.0001; **Fig. 2c, e and Supplementary Table 4**).

### Diverse denitrifiers across multiple phyla exhibit considerable structural diversity

From a cold seep genome catalog of 3,164 metagenome-assembled genomes (MAGs), we identified 142 *nosZ* sequences, which were categorized into *nosZ* clade I (n = 52) and *nosZ* clade II (n = 90) based on phylogenetic and structural analyses (**Supplementary Fig. 10 and Supplementary Table 8**). The relative abundance of the MAGs that comprised *nosZ* genes ranged from 0.0001 to 0.0263% in the 165 sediment samples of cold seeps (**Supplementary Table 8**). Nitrous oxide reductase (N_2_OR) is a copper-dependent enzyme that assembles into tightly linked, head-to-tail homodimers of 130 kDa, incorporating both a mixed-valent Cu_A_ center and a unique, tetranuclear Cu_Z_ site^62^. This arrangement allows N_2_O to bind across the Cu_Z_ site bridging the two copper centers^63^. Structurally, both cold seep *nosZ* clades I and II N_2_ORs feature a Cu_Z_ active site within the N-terminal seven-bladed β-propeller domain and a Cu_A_ site in the C-terminal cupredoxin domain, without any transmembrane α-helices (**Supplementary Figs. 11-12**). Their N-terminal configurations differ to align with their physiological functions (**Fig. 3a**), adopting signal peptides^64^ for either the Tat pathway in clade I or the Sec pathway in clade II (**Supplementary Figs. 11-12 and Supplementary Table 14**). Additionally, some *nosZ* clade II variants possess a C-terminal α-helix (**Fig. 3a, Supplementary Figs. 11b, 12b**), potentially enhancing the stability of their active sites. Critically, the Cu_A_ and Cu_Z_ sites within each monomer are too far apart (40Å) for effective electron transfer, but adjacent monomers have their Cu_A_ and Cu_Z_ sites just 10Å apart, facilitating electron flow^62^. Consequently, the C-terminal α-helix of *nosZ* clade II surrounds the periphery of Cu_A_ and Cu_Z_ sites of adjacent monomers in the structure. We used all the above-described structural features to query the metagenomic datasets and detect *nosZ* genes with very high confidence and resolution.

**Figure 3.**
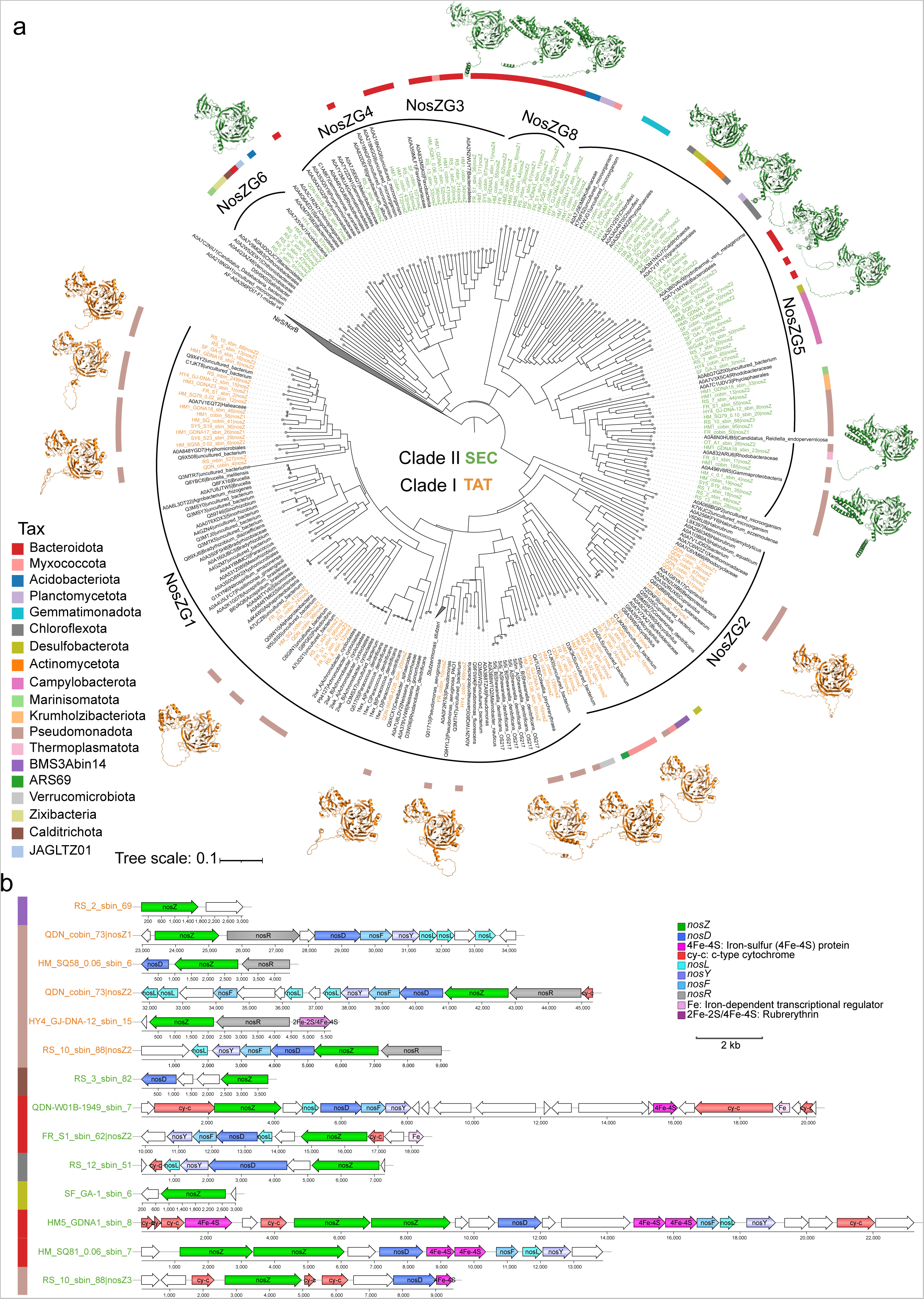
Structure trees of NosZ and *nos* clusters from genomes harboring Clade I or Clade II *nosZ* genes. (a) Structure trees created with Foldtree and affiliated taxonomic phylum of NosZ in MAGs. *nosZ* Clade I (NosZG1, NosZG2) is shown in orange, while *nosZ* Clade II (NosZG3, NosZG4, NosZG5, NosZG6, NosZG8) is shown in green. NosZG8, belonging to *nosZ* Clade II, was only found through structural analysis. Scale bar indicates the mean number of substitutions per site. Surrounding the tree, the affiliated taxonomic phylum and AlphaFold-predicted 3D structures of representative NosZ are displayed. (b) Comparison of *nos* clusters from genomes harboring Clade I or Clade II *nosZ* genes. The *nos* clusters of Clade II harbor the atypical *nosZ* and encode predicted iron-sulfur-binding proteins (labeled “4Fe-4S” or “2Fe-2S”) and *c*-type cytochromes (cy-*c*). Accessory genes (*nosD*, *nosF*, *nosL*, and *nosY*) are generally conserved across *no*s clusters with both typical (Clade I) and atypical *nosZ*. Non-colored genes in the operons have no orthologs in any other known *nos* cluster. *nosR* and *nosX* are associated exclusively with typical *nos* clusters. Details for the taxonomy and annotations of *nosZ*-containing contig are provided in **Supplementary Tables 8, 11-12**.

The distribution of the 142 *nosZ* genes across MAGs spans one archaeal and 18 bacterial phyla, reflecting the wide phylogenetic breadth of nitrous-oxide reducers in these environments (**Supplementary Fig. 10a and Supplementary Table 8**). The most common phyla containing *nosZ* genes are *Pseudomonadota* (n = 56) and *Bacteroidota* (n = 36), with fewer occurrences in *Campylobacterota*, *Myxococcota*, *Chloroflexota*, *Desulfobacterota*, *Gemmatimonadota* and other phyla (**Fig. 3a**). *NosZ* clade I genes are present in *Alphaproteobacteria*, *Gammaproteobacteria*, and a few other phyla, while clade II genes are more widespread, found in 15 bacterial groups and the archaeal phylum *Thermoplasmatota* (**Supplementary Fig. 10 and Supplementary Table 8**). This reveals six additional phyla (*Campylobacterota*, *Desulfobacterota*, *Krumholzibacteriota*, *Myxococcota*, *Planctomycetota* and *Zixibacteria*) capable of reducing nitrous oxide, considerably expanding the known genetic diversity of N_2_O reducers in deep-sea cold seep sediments. Most clade II MAGs contain multiple *nosZ* genes, with six MAGs containing two copies and one containing three copies, contrasting with the single *nosZ* gene typically found in clade I MAGs from cold seeps (**Fig. 3b, Supplementary Fig. 13 and Supplementary Table 12**). Notably, we identified a MAG, RS_10_sbin_88, from *Gammaproteobacteria* that carries one copy of both *nosZ* clade I and II genes. The *nosZ* clade I genes of *Pseudomonadota* were transcribed at high levels, up to 150.71 TPM, while *nosZ* clade II genes were transcribed at moderate to high levels in several phyla, up to 41.92–200.31 TPM, especially in *Campylobacterota* and *Pseudomonadota* (**Supplementary Table 9-10**).

The *nos* gene clusters, essential for N_2_OR maturation and function, show substantial differences between the *nosZ* clade I and clade II. The *nos* clusters of *nosZ* clade II contain more genes than those of clade I (**Fig. 3b and Supplementary Fig. 13**). Both clades share three ABC transporter complex genes (*nosD*, *nosF* and *nosY*) and a copper chaperone gene (*nosL*) necessary for assembling the Cu_Z_ center^65^, typical of denitrifier *nos* clusters of clade I. There is no evidence suggesting differences in the copper transport process from NosL to NosZ via NosD between the clades^51^. Unlike clade I, the *nos* clusters of clade II lack the *nosR* gene, typically associated with electron transfer to NosZ^51^. Instead, they feature three extra genes coding for a 4Fe-4S dicluster domain protein, an iron-dependent transcriptional regulator, and a *c*-type cytochrome oxidase (**Fig. 3b and Supplementary Fig. 13**), which are likely involved in electron transport and Sec pathway translocation for functional integration with the cytochrome *c* maturation system.

Additionally, from 3,164 MAGs, we identified five *nod* genes characterized by conserved enzymatic active center residues, distinguishing them from the related nitric oxide reductase (*nor*) by the substitution of “Thr” with “Ile”, “His” with “Asp”, and “Glu” with “Gln”^41^ (**Fig. 4**). The relative abundance of MAGs comprising *nod* genes ranged from 0.0004 to 0.0133% in the 165 samples of cold seeps (**Supplementary Table 15**). Structurally, the nitric oxide dismutase from cold seeps consistently exhibited conserved α-helix domains—four enzymes had 14 α-helices, and one had 13, reflecting notable structural conservation (**Supplementary Fig. 14a-e and Supplementary Table 16**). These *nod* genes are found in five MAGs across two bacterial phyla, indicating a broader presence of oxygenic denitrifiers than previously understood (**Fig. 4a-b and Supplementary Table 15**)^13, 34, 41^. Specifically, three of these MAGs are attributed to the *Planctomycetota* phylum (two from UBA1135 and one from *Scalinduaceae*), and two to the *Bacteroidota* phylum (as *Maribacter_A sp023141835* and *Cecembia rubra*from the orders *Flavobacteriales* and *Cytophagales*, respectively). This expanded diversity suggests more widespread bacterial capabilities for nitric oxide dismutation at cold seeps than previously appreciated through metagenomic or environmental studies^66^. The gene cluster for nitric oxide dismutase in UBA1135 is conserved across two genomes, featuring a helix-turn-helix domain and electron transport-associated proteins, including ferredoxin oxidoreductase and a 4Fe-4S dicluster domain (**Supplementary Table 17**). In *Cecembia rubra*, the *nod* gene cluster includes not only the standard *nos* cluster (*nosZ*, *nosD*, *nosF*, *nosY*, *nosL*) but also elements that regulate nitric oxide signaling and electron transport. However, in the other two MAGs from *Scalinduaceae* and *Maribacter_A* sp023141835, the genes downstream of *nod* are categorized as having domains of unknown function. Additionally, most of these *nod*-containing MAGs possess genes for other nitrogen metabolic processes (**Supplementary Table 15**) such as genes for hydroxylamine dehydrogenase (*hao*), nitric oxide reduction (*nor*), nitrous oxide reduction (*nosZ*), nitrate reduction (*nap*/*nir*), nitrite reduction to nitric oxide (*nir*) and nitrite reduction to ammonia (*nrf*/*nir*).

**Figure 4.**
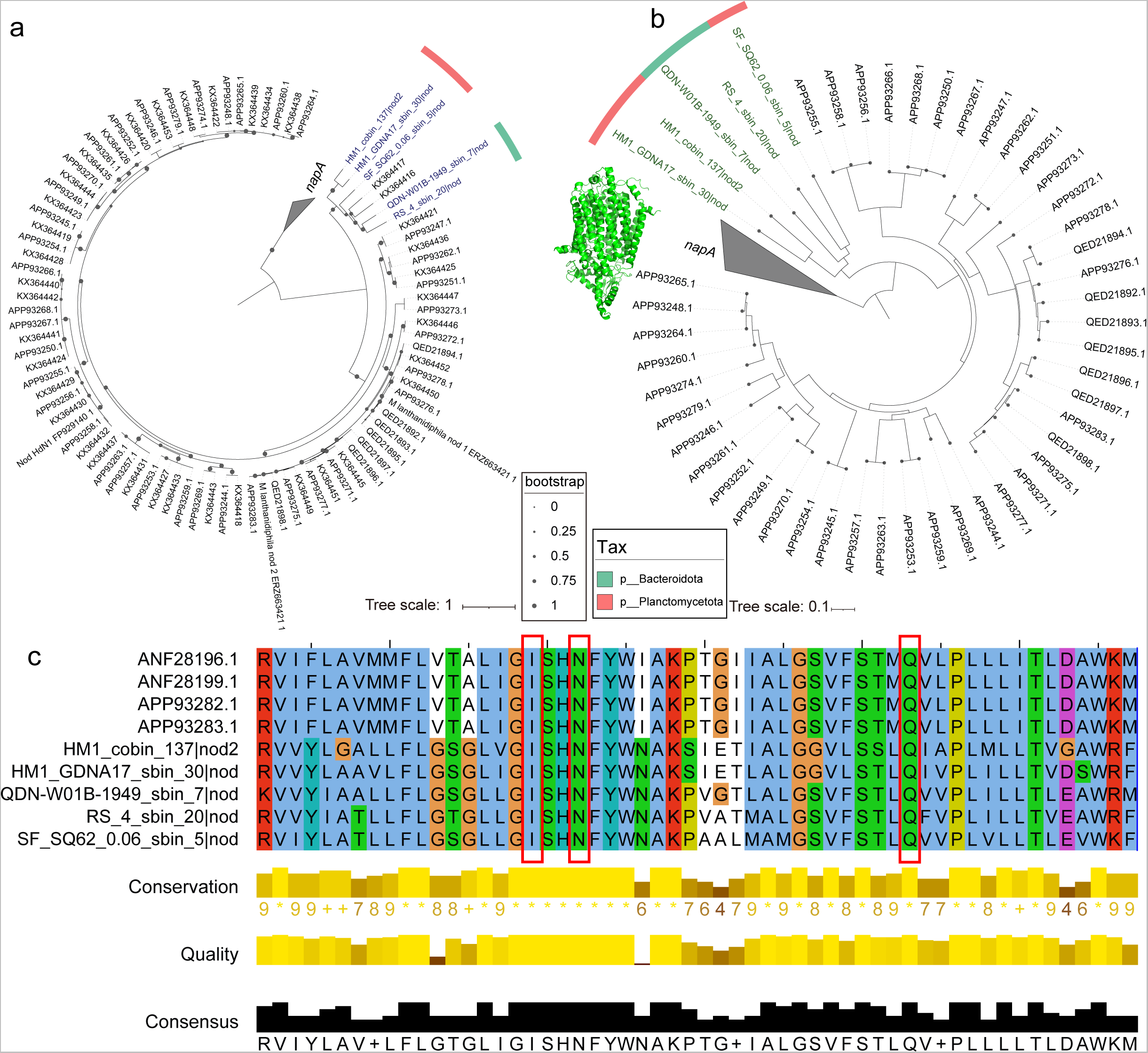
Trees and alignment of Nod recovered from 3,164 cold seep MAGs. (a) Maximum-likelihood phylogenetic tree of *nod* genes. (b) Structure tree of Nod created with Foldtree. Each Nod is labeled in dark blue and dark green in phylogenetic tree and structure tree, respectively. Scale bar indicates the mean number of substitutions per site. (c) Alignment of Nod and reference sequences. It shows the same diagnostic substitutions as the putative *nod* gene of the known NO-dismutating microbe *Methylomirabilis oxyfera*. Details for Nod MAGs are provided in **Supplementary Tables 15**.

### Anammox capabilities are found in multiple phyla beyond *Planctomycetota*

Further analyses of the 3,164 MAGs identified 265 *hzsA* genes, the diagnostic gene of bacteria capable of anammox (**Fig. 5a; Supplementary Table 18**). The relative abundances of these *hzsA-*containg MAGs ranged from 0.0002% to 0.0201% in the 165 samples from cold seeps (**Supplementary Table 18**). These *hzsA* gene sequences, characterized by a motif binding to a pentacoordinated *c*-type heme in the hydrazine synthase alpha subunit^52^, were distributed across 94 bacterial MAGs spanning 10 bacterial phyla (**Supplementary Fig. 15 and Supplementary Table 18**). This distribution extends beyond the previously recognized anammox bacteria of the *Brocadiales* order within the *Planctomycetota* phylum^42–44, 67^. These *hzsA*-containing MAGs are members of *Planctomycetota* (n = 55), *Bacteroidota* (n = 16), *Acidobacteriota* (n = 10), *Verrucomicrobiota* (n = 6), *Sumerlaeota* (n = 1), *JABMQX01* (n = 1), *Calditrichota* (n = 1), *Desulfobacterota* (n = 1), *Fibrobacterota* (n = 1), *Gemmatimonadota* (n = 1) and *Zixibacteria* (n = 1) (**Supplementary Fig. 15 and Supplementary Table 18**). The *hzsA* genes of *Acidobacteriota*, *Bacteroidota*, *Calditrichota*, *Planctomycetota* and *Sumerlaeota* were transcribed at moderate to high levels, up to 56.75–414.23 TPM, whereas fewer transcripts from *Verrucomicrobiota* were detected (**Supplementary Tables 19**).

**Figure 5.**
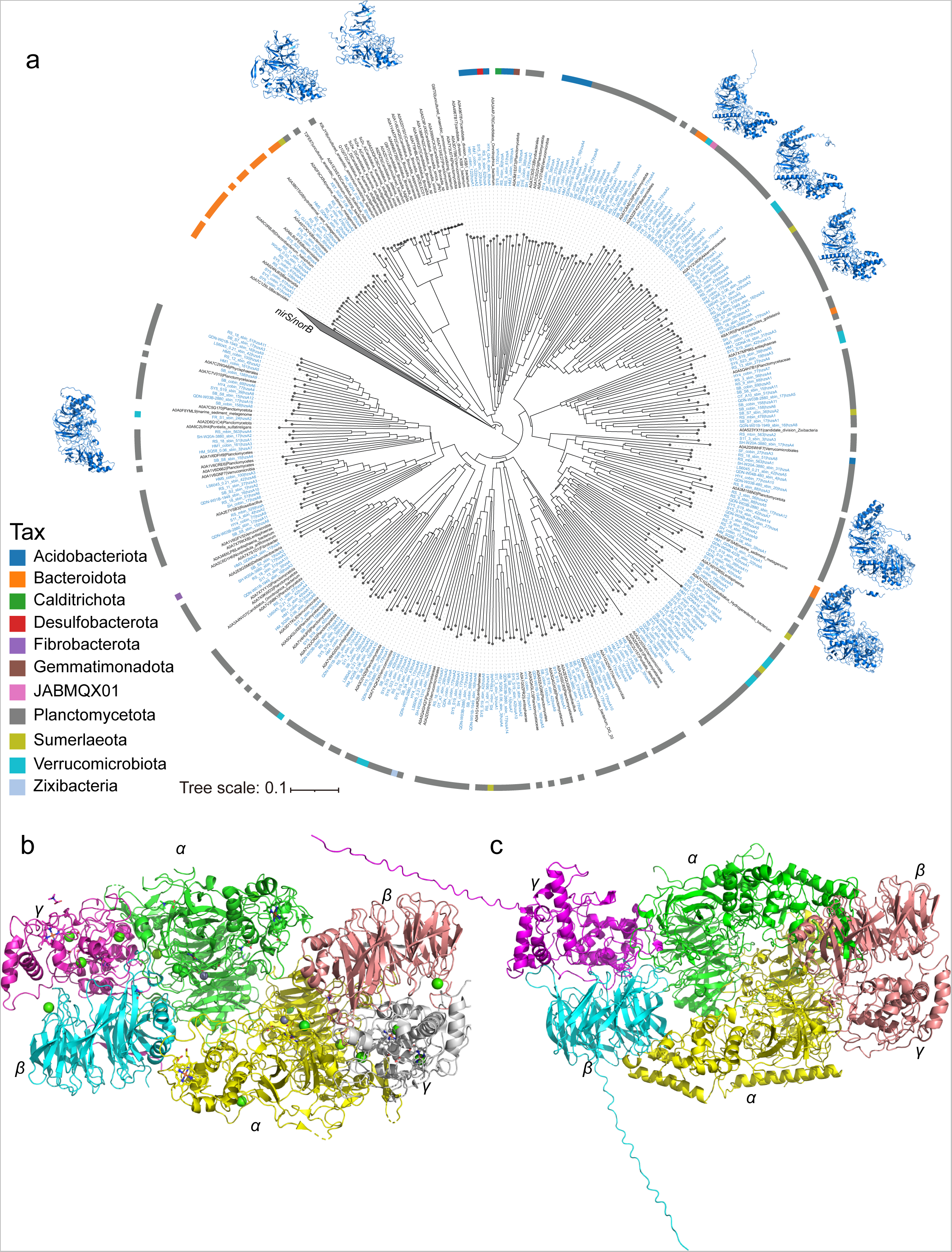
Structure tree of HzsA recovered from 3,164 cold seep MAGs created with Foldtree and predicted structure of HZS. (a) Structure tree created with Foldtree and affiliated taxonomic phylum of HzsA in MAGs. Each cold seep HzsA is labeled in blue in the structure tree. Scale bar indicates the mean number of substitutions per site. Surrounding the tree, the affiliated taxonomic phylum and AlphaFold-predicted 3D structures of representative HzsA are displayed. (b) Reference structure of HZS in 5C2V belonging to *Candidatus* Kuenenia stuttgartiensis (*Planctomycetota*) and (c) FR_S1_sbin_24 belonging to *Verrucomicrobiales* (*Verrucomicrobiota*). In the HZS complex structure: α-subunits are colored green and yellow, β-subunits are colored blue and pink, and γ-subunits are colored grey and magenta. Details for the taxonomy of *hzsA*-containing contig are provided in **Supplementary Tables 18**.

Among the 94 MAGs, 27 contain 42 *hzd* genes, which encode hydrazine dehydrogenase for oxidizing N_2_H_4_ to N_2_ (**Supplementary Table 21**). Notably, only two belong to the *Ca.* Scalinduaceae family within the *Planctomycetota* phylum, and both do not contain the complete *hzsBC* genes (**Fig. 6 and Supplementary Table 22**). However, 37 MAGs do contain these genes (**Supplementary Tables 22-23**), forming a multienzyme complex HZS-αβγ^68^. Of these, five MAGs comprise multiple *hzsABC* clusters and are associated with *Phycisphaerae* (n = 3), *Verrucomicrobiae* (n = 1), and *Bacteroidia* (n = 1). These findings considerably broaden the recognized phylogenetic diversity and environmental presence of anammox bacteria.

**Figure 6.**
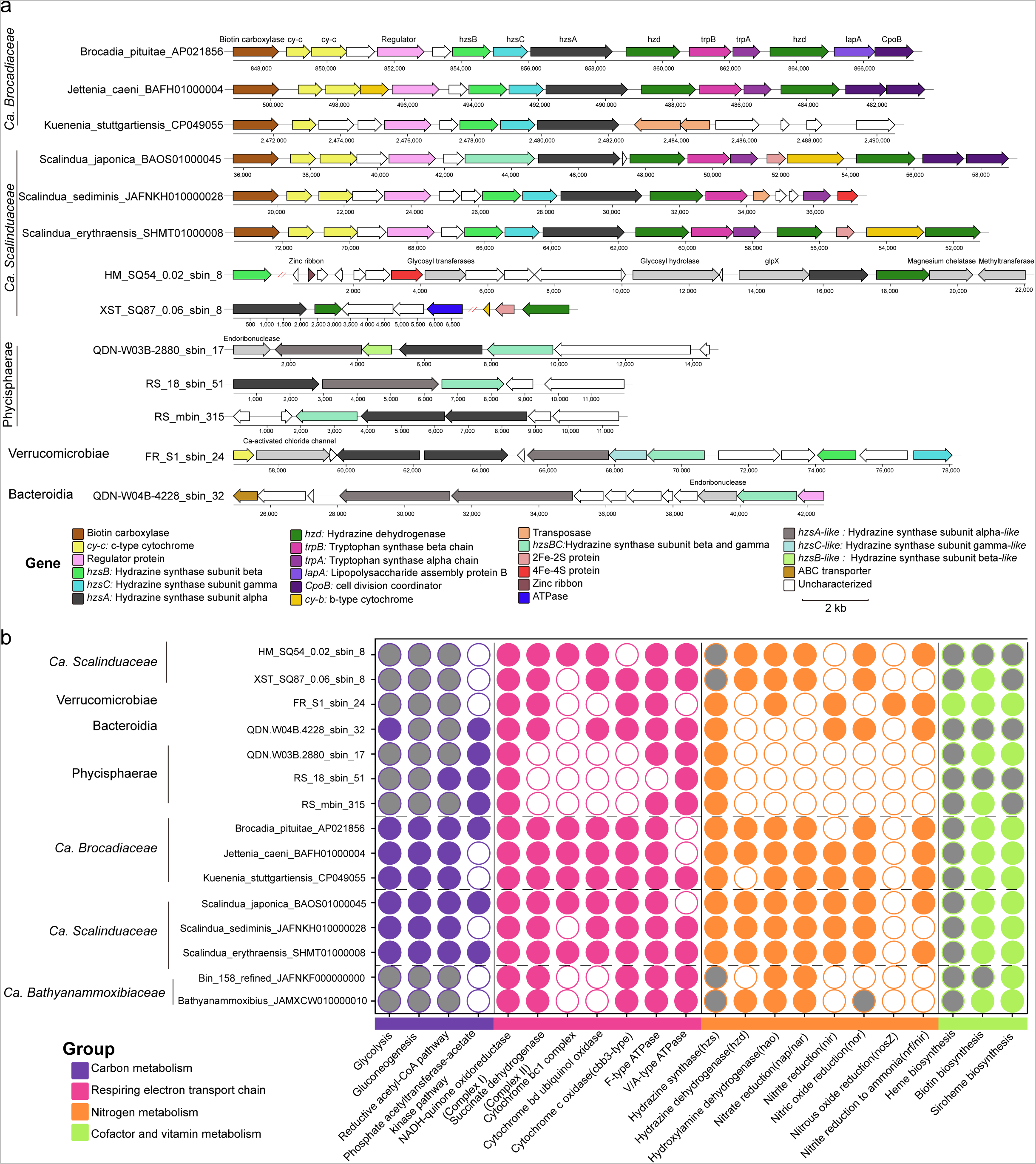
Comparison of *hzs* gene clusters and metabolic features from traditional and novel genomes harboring *hzsABC* genes. (a) Conservation of key genes encoding hydrazine metabolism in anammox bacteria genomes. (b) Comparison of metabolic potential of anammox bacteria. Filled circles represent the presence of genes encoding the metabolic process, grey circles denote partial presence, and open circles indicate absence. Details for *hzsA*-containing annotations are provided in **Supplementary Tables 20-23**.

We then carefully examined the five MAGs with multiple *hzsABC* clusters and two *Scalinduaceae* MAGs, confirming they were not contaminated during binning (**Supplementary Table 24**). We found that the MAGs not only shared a similar gene cluster arrangement (**Fig. 6a; Supplementary Table 22**) but also structural similarities with known anammox bacteria (**Fig. 5a**). In addition to the essential genetic machinery for anammox metabolism, including hydrazine synthase (*hzs*) for converting NO and NH_4_^+^ to N_2_H_4_, and hydrazine dehydrogenase (*hzd*) for oxidizing N_2_H_4_ to N_2_, the MAGs also possess additional nitrogen metabolic processes such as genes for nitrate reduction (*nap*/*nir*), nitrite reduction to nitric oxide (*nir*) and nitrite reduction to ammonia (*nrf*/*nir*)^69^ (**Fig. 6b and Supplementary Table 25**). Notably, we identified a MAG affiliating with *Verrucomicrobiae*, FR_S1_sbin_24, which contains at least two copies of the *hzsA*, *hzsB*, and *hzsC* genes in a single contig (**Fig. 6a and Supplementary Table 22**), interspersed with several unknown proteins and upstream genes related to *c*-type cytochromes and Ca-activated chloride channels. This MAG also encodes both hydroxylamine oxidoreductase (Hao) for nitrite reduction^70^ and NosZ (**Fig. 6b**), suggesting a versatile metabolic capability.

Both traditional and novel HzsA proteins encoded in the seven MAGs are structured into three distinct domains (**Fig. 5a, Supplementary Figs. 16a, b, and 17a, b**): a six- bladed β-propeller N-terminal domain, a middle domain that binds a pentacoordinated *c*-type heme (heme αI), and a C-terminal domain featuring a bis-histidine-coordinated *c*-type heme (heme αII)^52^. The heme αI substantially differs from typical heme *c* sites by coordinating a zinc ion. Traditional HzsA proteins lack an N-terminal transmembrane α-helix (**Supplementary Fig. 16c, d**), as do the novel HzsA proteins described here (**Supplementary Fig. 17c, d**). However, only the novel HzsA proteins include N-terminal signal peptides indicative of the Sec secretion pathway (**Supplementary Fig. 17e, f**). Additionally, these proteins exhibit significant variation in their C-terminal structures, particularly the novel HzsA proteins, which include a C-terminal α-helix that may stabilize the binding at the heme αI active center (**Fig. 5a and Supplementary Figs. 16a, b, 17a, b**). These proteins assemble into one or two heterotrimer complexes as predicted by AlphaFold Multimer (**Fig. 5c, Supplementary Fig. 22**). In each complex, one of the β and γ subunits of the novel HzsA are fused into a single polypeptide chain, consistent with earlier studies^71^. The novel HzsA proteins retain essential domains for heme binding and electron transfer, indicating they are functionally capable of catalyzing anammox reactions. Structural adaptations, such as N-terminal signal peptides and C-terminal α-helices, optimize these proteins for the unique conditions of cold seeps, showcasing the evolutionary adaptability of anammox bacteria in nitrogen cycling.

### Conclusions

In the past, gene annotations primarily relied on sequence similarity, which often overlooked genes that are distantly related yet functionally similar. However, over long evolutionary time scales, multiple substitutions at the same site can cause uncertainty in sequence alignment. Structures evolve at a slower rate than the underlying sequence mutations and are more conserved, emphasizing the importance of protein structural analysis^72^. By employing protein structural similarities and phylogenetic analysis, we have discovered that the NosZ clade II is notably more diverse and abundant in deep-sea cold seeps than previously assumed. This diversity may have significant implications for the role of NosZ clade II in N_2_O consumption in cold seeps. Members of the *Planctomycetota* phylum, as well as the orders *Flavobacteriales* and *Cytophagales*, might be notable contributors to nitrogen loss through nitric oxide dismutation. These organisms might also act as oxygen producers which could be linked to the aerobic methane and sulfide oxidation within anoxic layers of cold seep ecosystems^73^. Crucially, our findings also indicate that anammox bacteria are found in multiple phyla (e.g. *Bacteroidota*, *Acidobacteriota*, and *Verrucomicrobiota*) beyond *Planctomycetota*, and that these overlooked lineages actively express anammox genes. Our findings highlight the importance of nitrogen loss through denitrification and anammox processes at cold seeps. Overall, this study provides evidence supporting the presence of numerous novel, cold-adapted microbial lineages involved in denitrification (including oxygenic denitrification) and anammox processes. It establishes cold seeps as considerable nitrogen-loss hotspots in the deep sea and as important contributors to the global nitrogen cycle, broadening the recognized roles of cold seeps beyond serving as oases of diversity, productivity, and methane removal.

## Materials and methods

### Sampling and geochemical measurements

A total of 301 samples from 33 push cores were collected by a remotely operated vehicle (ROV) at Lingshui, Haima and Site F cold seeps in the South China Sea between 2020 and 2023 (**Supplementary Fig. 1**). Upon retrieval on deck, sediment cores were immediately placed in a helium-filled glove bag. Porewater was extracted at 2-cm depth intervals using Rhizon samplers with 0.2 μm pore size (Rhizosphere, Netherlands). Collected porewater samples for used for metal analysis were acidified to pH ∼2 with HNO_3_ (Optima grade, Thermo Scientific, USA), and stored at 4 °C before analysis. For nitrogen gas measurement in sediments, 3 mL of sediments at 2- cm intervals was transferred using a 5 mL cut-off syringe into a 22 mL serum vial, which was then crimp-sealed and stored at 4 °C. Samples of porewater and sediment for analyzing nutrients, sulfate, and calcium were stored at −20 °C until analysis.

The concentrations of N_2_ along with its δ^15^N values in the headspace gas were measured using gas chromatography with thermal conductivity detection (Agilent, USA) and a continuous-flow isotope-ratio mass spectrometer (SerCon, UK). The concentrations of ammonium (NH_4_^+^), nitrite (NO_2_^−^), and nitrate (NO_3_^−^) in porewater were determined using a Quattro continuous flow analyzer (SEAL Analytical AA3, Germany). Sulfate (SO_4_^2–^) and calcium (Ca^2+^) concentrations were determined using a Dionex Ion Chromatograph (Thermo Scientific Dionex, USA). Dissolved metals (Fe^2+^, Cu^2+^ and Zn^2+^) in porewater were measured using ICP-MS (Thermo Scientific, USA). Sediment total organic carbon (TOC) and total nitrogen (TN) were measured using a Vario Micro Cube elemental analyzer (Elementar, Germany), after the sediments were treated with 1 M HCl to remove carbonates.

### Determination of nitrogen-loss rates

Nitrogen-loss rates were determined using 37 samples from 8 push cores (Lingshui-10/-11, Haima-3/-4/-5/-6/-7/-8) collected by an ROV during 2023 from the cold seeps of Lingshui and Haima, with water depths ranging from 1350 to 1822 meters. Additionally, 11 samples from 3 push cores (Shenhu-1/-2/-3) were collected from the non-seep area in Shenhu, at depths of 489 to 1457 meters in the South China Sea (**Supplementary Fig. 1**). Potential nitrogen-loss rates were measured using N isotope-tracing techniques as follows^74, 75^.

Briefly, slurries were prepared using collected sediments and artificial seawater matching *in situ* salinity, at a sediment/water volume ratio of 1:7. The mixture was purged with helium for 30 min and stirred vigorously to ensure homogeneity. Gas-tight borosilicate vials (Labco Exetainers) were then filled with slurries under a helium atmosphere. Subsequently, the vials were pre-incubated at near *in situ* temperature (4 °C) for 48 h to remove residual NO_x_ ^−^ (NO_3_ ^−^ + NO_2_ ^−^) and dissolved oxygen. The slurries were divided into three groups, each amended with different nitrogen compounds in helium-purged stock solutions: (1) ^15^NH_4_^+^ (99.12%), (2) ^15^NH_4_^+^ + ^14^NO_3_^−^, and (3) ^15^NO_3_ ^−^ (99.21%). The final concentration of ^15^N compounds in each vial was 75 μM. To halt the incubation, 200 μL of 50% ZnCl_2_ solution was injected into each vial. The ^15^N labeled N_2_ were measured using a membrane inlet mass spectrometer (MIMS, HPR-20, Hiden Analytical, UK). Denitrification and anammox rates were estimated from the accumulation of ^29^N_2_ and ^30^N_2_ during the slurry incubation^16, 74^. The respective contributions of denitrification and anammox to ^29^N_2_ production were quantified using equation (1).

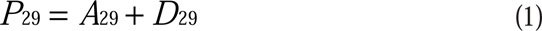

where *P_29_*(nmol cm^-3^ h^-1^) denotes the total ^29^N_2_ production rates, *D_29_* (nmol cm^-3^ h^-1^) and *A_29_* (nmol cm^-3^ h^-1^) denote the production rates of ^29^N_2_ from denitrification and anammox, respectively. Here, *D_29_* was obtained by equation (2), assuming random paring of ^14^N and ^15^N from ^14^NO_3_^−^ or ^15^NO_3_^− 76, 77^.

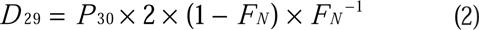

where *P_30_*(nmol cm^-3^ h^-1^) denotes the total ^30^N_2_ production rates, *F_N_* (%) denotes the fraction of ^15^N in NO_3_^−^, which was obtained from the added ^15^NO_3_ ^−^ and the measured residual ambient NO_x_ ^−^, ranging from 97.8% to 99.7%. The potential rates of denitrification and anammox were quantified by equations (3) and (4).

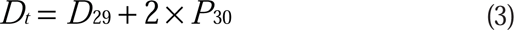

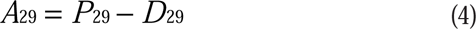

where *D_t_* and *A_29_* (nmol cm^-3^ h^-1^) denote the denitrification and anammox rates, respectively. By convention, the percent of N_2_ production accounted for anammox is abbreviated as *ra* (%).

### Metagenomic data processing

The metagenomic datasets from 165 samples were collected from 16 cold seep sites worldwide, including oil and gas seeps, methane seeps, gas hydrates, asphalt volcanoes, and mud volcanoes (**Supplementary Fig. 1**). Non-redundant gene and genome catalogs were constructed as described in our previous study^59^. Briefly, metagenomic sequence data were quality controlled and assembled into contigs. Protein-coding sequences were then predicted and clustered to create a non-redundant gene catalog consisting of 147,289,169 representative clusters. Salmon (v1.10.2)^78^ was used to calculate gene abundance in each metagenome, which was then normalized to genes per million (GPM). Functional annotations were performed using eggNOG-mapper (v2.1.9) with default parameters^79, 80^.

Contigs longer than 1000 bp were selected for subsequent binning, and the produced metagenome-assembled genomes (MAGs) underwent dereplication at 95% average nucleotide identity^59^.A total of 3,164 representative MAGs were obtained. The relative abundance of each MAG was calculated using CoverM (v0.6.1; https://github.com/wwood/CoverM; parameters: -m relative_abundance --trim-min 0.10 --trim-max 0.90 --min-read-percent-identity 0.95 --min-read-aligned-percent 0.75). The taxonomy of each MAG was assigned using GTDB-Tk v2.1.1 with reference to GTDB R207 database^81^.

### Identification of nitrous oxide reductase gene (*nosZ*)

For *nosZ* gene database search (see workflow in **Supplementary Fig. 4a**), we first queried protein sequences from the non-redundant gene catalog and 3,164 MAGs against NCycDB^82^ using DIAMOND (version 2.0.14)^83^ in blastp mode (-k 1 -e 0.0001 -p 5). Subsequently, the *nosZ* reference sequences (n = 403) from the Greening lab metabolic marker gene databases^84^ were used to search for potential *nosZ* sequences in the non-redundant gene catalog and 3,164 MAGs with DIAMOND blastp (version 2.0.14; --id 50)^83^. Additionally, hidden Markoc models (HMMs) of *nosZ* clade I (TAT-dependent nitrous-oxide reductase) and *nosZ* clade II (Sec-dependent nitrous-oxide reductase) were obtained from NCBI’s Protein Family Models using accession “TIGR04244.1” ^85^ and “TIGR04246.1” ^86^, respectively. These HMMs were used to screen proteins from the non-redundant gene catalog and 3,164 MAGs with hmmsearch in HMMER v3.3.2 using the parameter -E 1e-5. All *nosZ* genes identified with the above three methods were merged, and any sequences shorter than 400 amino acids were excluded.

The phylogenetic trees of the amino acid sequences of filtered genes were constructed to validate the phylogenetic clades of *nosZ* against reference sequences. Sequences were aligned using MUSCLE (v3.8.1551)^87^ and trimmed with TrimAL (v1.4.1)^88^ with default settings. Maximum-likelihood trees were constructed with IQ-TREE (v2.2.0.3)^89^ with the “-m MFP -B 1000” options. The produced tree was visualized and beautified using Interactive tree of life (iTOL; v6)^90^. Meanwhile, ESMFold ^91^ was applied to predict the structure for each filtered gene. The 154 reference protein structures of NosZ were downloaded from AlphaFoldDB^92^ and Protein Data Bank (PDB)^93^. A structural tree of NosZ was constructed using Foldtree (https://github.com/DessimozLab/fold_tree) based on a local structural alphabet^94^ and visualized using iTOL (v6)^90^. The structure of each gene in both the phylogenetic and structural trees was predicted using AlphaFold2 ^95^ and aligned against PDB using Foldseek (v8.ef4e960)^96^ with parameters “--tmscore-threshold 0.5 -e 0.001”. Cupredoxin-related protein active domains of nitrous-oxide reductase were investigated against the ECOD (Evolutionary Classification Of protein Domains) database^97^ with a TM-score > 0.5 using Foldseek easy-search module^96^.

### Identification of nitric oxide dismutase gene (*nod*)

For *nod* gene database search (see workflow in **Supplementary Fig. 4b**), we initially downloaded reference protein sequences (n = 1036) in NCBI’s databases to build an HMM model (available at https://doi.org/10.6084/m9.figshare.25650927). Then, *nod* genes in the non-redundant gene catalog and 3,164 MAGs were extracted using the above HMM model with hmmsearch in HMMER v3.3.2 using “-E 1e-5”. All potential *nod* genes shorter than 200 amino acids were excluded from further analysis. Following the methodology used for *nosZ* genes, these *nod* genes underwent verification through phylogenetic and structural trees. The reference protein structures for 44 Nod proteins were predicted using AlphaFold2 ^95^. The structures of the filtered *nod* genes were then predicted using AlphaFold2 and aligned with the reference protein structures using Foldseek (v8.ef4e960)^96^. Additionally, diagnostic amino acid residues in the active center of the enzyme^41^ were identified in all proteins with a TM- score > 0.5 using MAFFT (EMBL-EBI)^98^ and visualized with Jalview^99^.

### Identification of hydrazine synthase and hydrazine dehydrogenase

To identify *hzsA* genes (see workflow in **Supplementary Fig. 4c**), protein sequences in the non-redundant gene catalog and 3,164 MAGs were firstly searched against NCycDB^82^, with the program DIAMOND blastp (version 2.0.14) ^83^. Then, reference *hzsA* sequences (n = 14) from the Greening lab metabolic marker gene databases^84^ were used to searched for potential *hzsA* sequences in the non-redundant gene catalog and 3,164 MAGs with DIAMOND blastp (version 2.0.14; --id 50)^83^. Additionally, *hzsA* genes of non-redundant gene catalog and 3,164 MAGs were extracted using HMMER v3.3.2 with the HMM profile “PF13486” from the InterPro database ^52^. All identified *hzsA* genes were merged, and sequences shorter than 400 amino acids were filtered out. These genes were then verified by constructing phylogenetic and structural trees. The structure of each gene was predicted using AlphaFold2^95^ and aligned with PDB^93^ using Foldseek (v8.ef4e960)^96^. Additionally, the active domain of the hydrazine synthase alpha subunit was searched in protein structures with a TM- score > 0.5 using Foldseek’s easy-search module^96^.

To ensure the accuracy of identifying MAGs with *hzsA* genes, MAGpurify (v2.1.2) was used to detect contamination in MAGs through a combination of features and algorithms including phylo-markers, clade-markers, tetra-freq, GC-content, and known-contam^100^. Additionally, *hzsB* and *hzsC* sequences in MAGs were extracted with HMMER v3.3.2 using the “nitro.cycle.sub.hmm” model from the Metascan metabolic HMM database^101^. To further refine our search, the protein complexes of the novel clade hydrazine synthase (HzsABC) were predicted using AlphaFold (v2.0; model_preset = multimer)^95^. All structures were visualized and exported as images using PyMOL (http://www.pymol.org)^102^. Additionally, the *hzd* gene associated with hydrazine dehydrogenase was identified in MAGs containing *hzsA* genes. To begin, we searched for reference protein sequences (n = 49) in NCBI’s databases to construct the HMM model (available at https://doi.org/10.6084/m9.figshare.25650927). Subsequently, *hzd* genes were extracted using hmmsearch in HMMER, applying the “-E 1e-5” parameter. The structures were then predicted using AlphaFold2 and aligned with reference proteins in the PDB databases using Foldseek (v8.ef4e960) with settings “--tmscore-threshold 0.5 -e 0.001”.

### MAG annotations and topological structure predictions

The MAGs were annotated using DRAM (v1.3.5)^103^ and Prokka (v1.14.6)^104^ with the default settings against KEGG, Pfam, MEROPS and dbCAN databases. Gene context was visualized using Chiplot (https://www.chiplot.online/), with the files produced by DRAM and Prokka as the input. DeepTMHMM (v1.0.24)^105^ and SignalP (v6.0)^106^ were employed to predict transmembrane topology and signal peptides of NosZ, Nod and HzsA proteins, respectively.

### Identification of *nifH*, *dsrA*, *mcrA* and mobile genetic elements

To identify *nifH* genes associated with nitrogen fixation, protein sequences in the non-redundant gene catalog was first searched against NCycDB^82^, using the program DIAMOND (version 2.0.14) ^83^ as mentioned above. Then, we utilized *nifH* reference sequences (n = 1271) from the Greening lab metabolic marker gene databases^84^ to search for potential *nifH* sequences in the non-redundant gene catalog using DIAMOND blastp (version 2.0.14; --id 50)^83^. Subsequently, *nifH* genes were extracted using the NCBI’s Protein Family Models with HMM accession “TIGR01287.1” by HMMER v3.3.2. Subsequently, *nifH* genes were extracted using NCBI’s Protein Family Models with the HMM accession “TIGR01287.1” via HMMER v3.3.2. The genes were then verified through phylogenetic analysis as described in our previous study^6^. Conserved motifs (CXXR)^6^ were analyzed using MAFFT(EMBL-EBI)^98^ and visualized with Jalview^99^.

For the identification of *dsrA* and *mcrA* genes, associated with sulfate reduction and anaerobic methane oxidation respectively, we employed eggNOG-mapper (v2.1.9; default parameters) for initial detection^79, 80^. The reductive *dsrA* genes were further detected using DiSco (v1.0.0)^107^, while oxidative *mcrA* genes were confirmed via phylogenetic analysis as described in our previous study^59^. Classification of contigs belonging to mobile genetic elements (plasmids, proviruses and viruses) was performed using Genomad v.1.5.0 with default parameters^108^.

### Transcriptional activities of nitrogen-loss genes

A total of 33 samples from various cold seeps—Haima, Qiongdongnan Basin, Shenhu area, and Jiaolong—were analyzed for metatranscriptomes, as outlined in our previous study^109^. Briefly, raw reads were quality filtered (parameters: --skip- bmtagger) using Read_QC module within the metaWRAP (v1.3.2) pipeline^110^. SortMeRNA (v2.1)^111^ was employed to remove ribosomal RNA from quality-controlled reads using default settings. The abundances of transcripts for *nosZ, nod* and *hzsA* genes were quantified by mapping filtered reads to a non-redundant gene catalog and genes of MAGs using Salmon (v.1.9.0; parameters: -validateMappings - meta)^78^. The results were expressed in transcripts per million (TPM).

### Statistical analyses

Statistical analyses were performed using R v4.2.3. The normality of the data was assessed using Shapiro–Wilk tests prior to further analyses. The Kruskal-Wallis rank-sum test was used to compare the abundances of nitrogen-loss genes across different depths, and their rates across different types of environments. Student’s *t*-tests were utilized to examine differences in the relative abundance of *nosZ*, *nod*, and *hzsA* genes at varying depths, as well as nitrogen-loss rates across non-seep and cold seep environments. Pairwise comparisons of environmental factors were conducted using Pearson’s correlation coefficients. Additionally, relationships between the relative abundance of *nosZ*, *nod* and *hzsA* genes and environmental factors was analyzed using Mantel tests.

## Data availability

The non-redundant gene catalog and the metagenome-assembled genomes (MAGs) catalog can be accessed at https://doi.org/10.6084/m9.figshare.22568107. The MAGs containing *nosZ*, *nod*, and *hzsA* genes, along with the protein structures and phylogenetic trees for NosZ, Nod, and HzsA, as well as all the HMM models developed in this study, are available at https://doi.org/10.6084/m9.figshare.25650927. All additional data supporting the findings of this study are provided within the article and its Supplementary Information Files.

## Code availability

The present study did not generate codes, and mentioned tools used for the data analysis were applied with default parameters unless specified otherwise.

## Acknowledgements

The work was supported by National Science Foundation of China (No. 92351304, No. 42376115 and No. 42030407), Natural Science Foundation Project of Xiamen City (No. 3502Z202373076), Natural Science Foundation of Fujian Province (No. 2023J06042), Scientific Research Foundation of Third Institute of Oceanography, MNR (No. 2022025 and No. 2023022). SER was supported by the Simons Foundation (824763) and funds from the Human Frontier Science Program (RGEC34/2023). We thank Chengpeng Li and Xinyue Liu for assistance in determining geochemical parameters, Weichao Wu for providing sediment samples, and Chris Greening for helpful discussions. We also express our gratitude to the captains, crews, and pilots of the *R/V KEXUE* as well as the ROV *Faxian* operation team for their support in collecting the samples.

## Author contributions

XD and QJ designed this study. QJ and ZZ performed the omics analysis. LC, JP and MW analyzed environmental factors. XL analyzed nitrogen-loss rates. YH contributed to discussions and methodology. SL, RZ, XZ, SER and BZ participated in discussions and data interpretations. LC, MW and JL collected cold seep sediment samples. QJ, XL, LC, and XD wrote the paper, with input from other authors.

