## Supplementary Figures 1-18 for "Cold seeps are hotspots of deep-sea nitrogen-loss driven by microorganisms across 21 phyla"

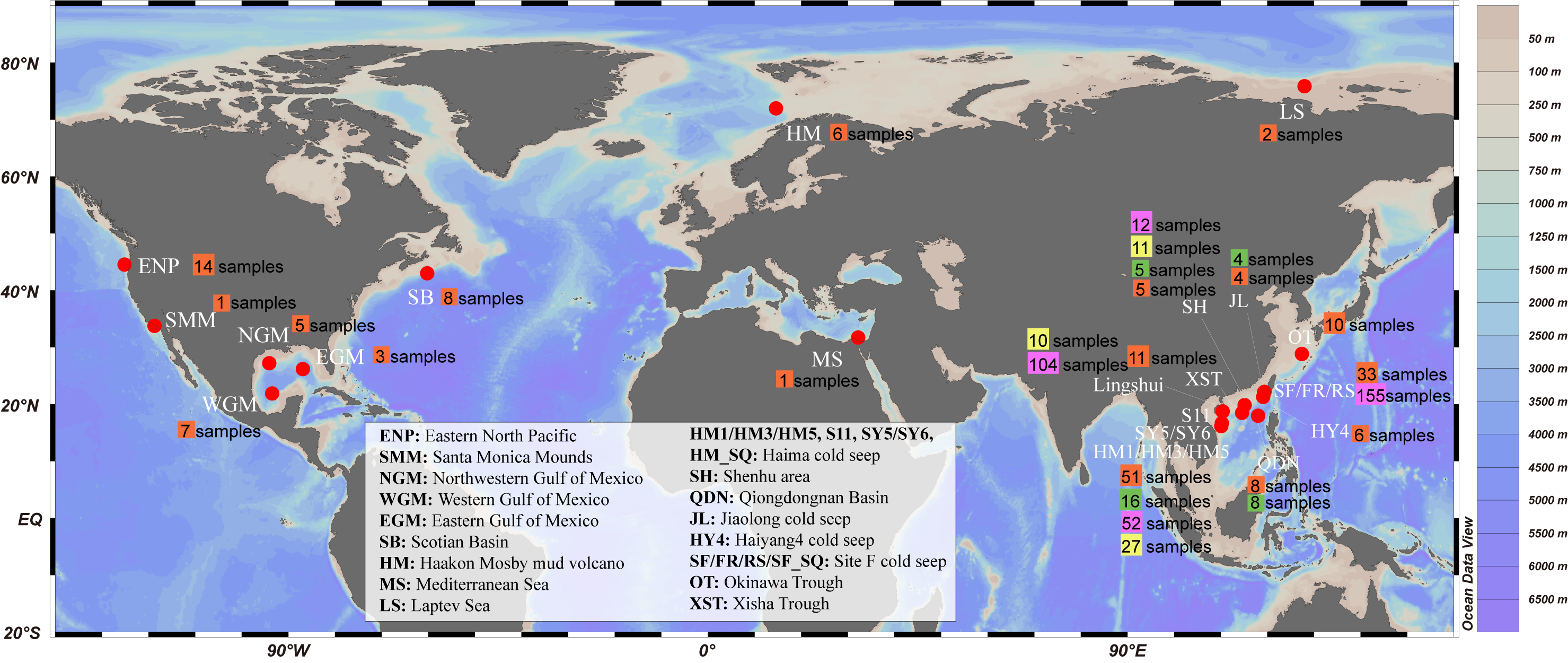

**Supplementary** Figure 1**. Location of the cold seeps study area**. Geographic distribution of cold seep sites for geochemistry analysis, nitrogen-loss rates analysis, metagenomic and metatranscriptomic data was indicated. Sites with pink squares denote that samples for geochemistry analysis were collected. Sites with yellow squares denote that samples for nitrogen-loss rates analysis were collected. Sites with orange squares denote that metagenomic data were collected, and sites with green squares denote that metatranscriptomic data were collected. The world map was drawn using the Ocean Data View (v5.1.7).

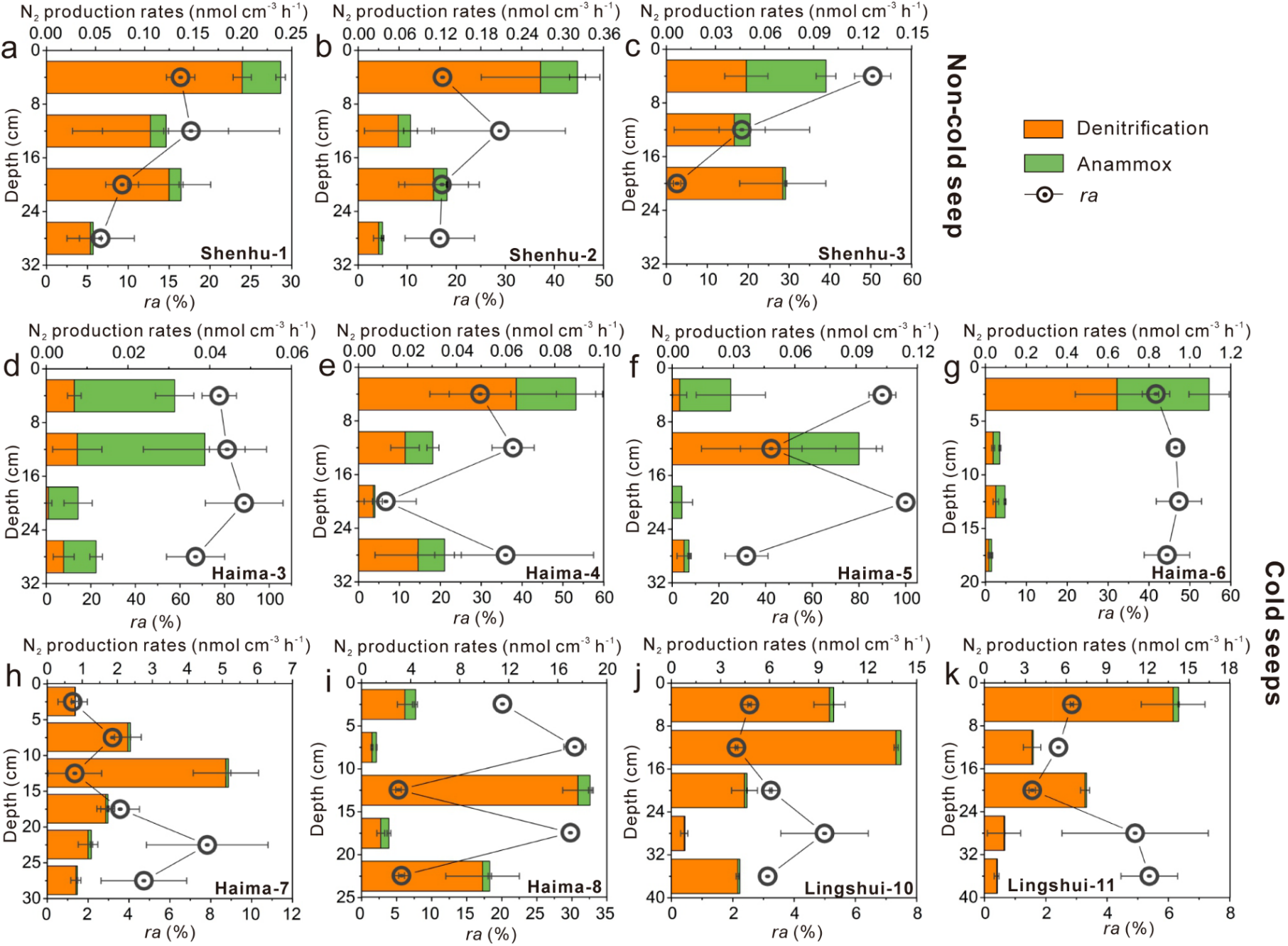

**Supplementary** Figure 2**.** **Vertical distributions of potential N_2_ production rates (denitrification, anammox) and the percent of N_2_ production as anammox (*ra*, %) in sediments of the non-seeps (a−c) and cold seeps (d−k)**. The errors denote standard deviation. Detailed data on rates profiles can be found in **Supplementary Table 2.**

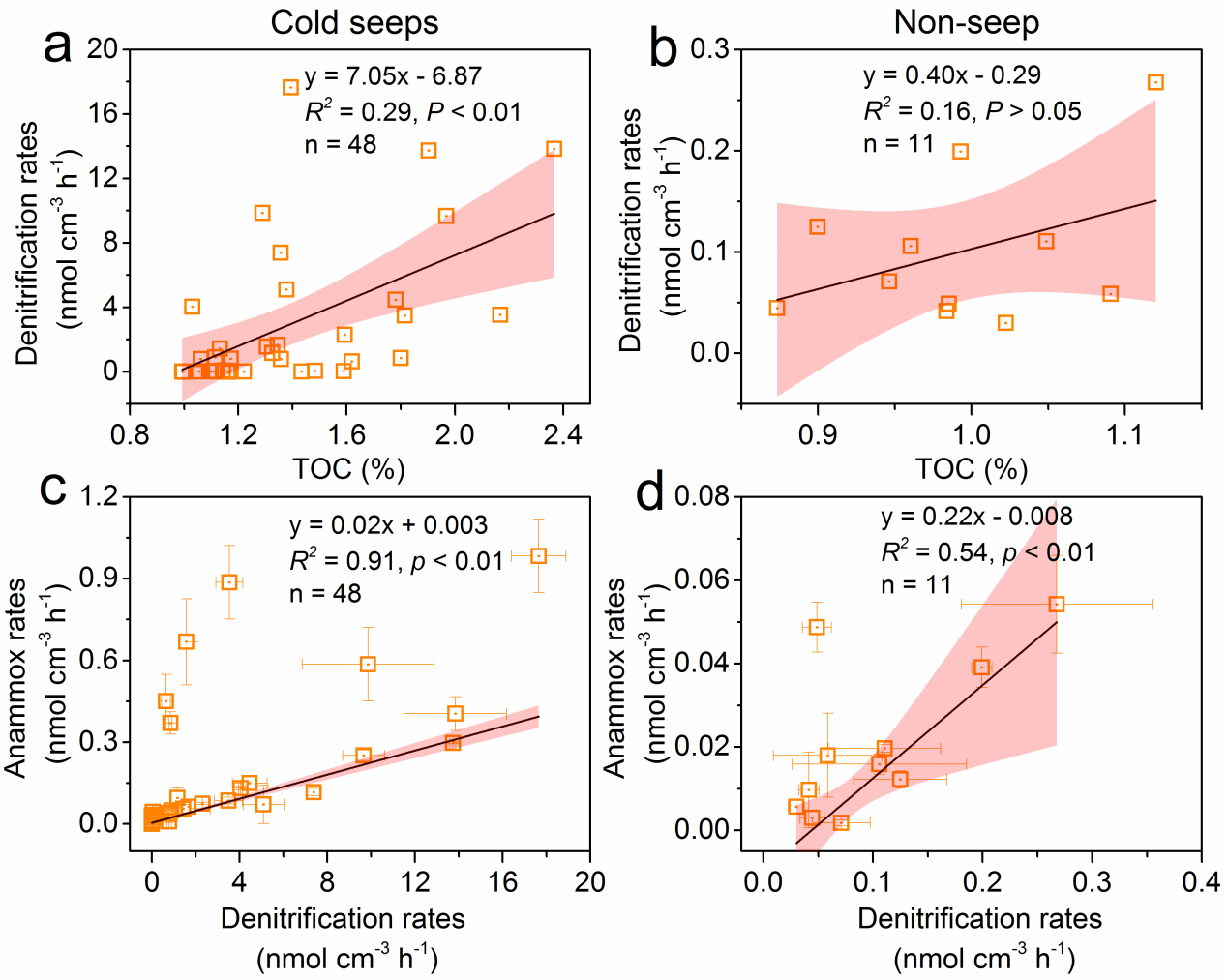

**Supplementary** Figure 3**.** **The relationships between measured denitrification rates and sediment TOC, as well as relationships between measured denitrification and anammox rates in sediments of cold seeps (a, c) and non-seep (b, d)**. Pink band indicates 95% confidence interval. Detailed data on rates profiles can be found in **Supplementary Tables 1 and 3.**

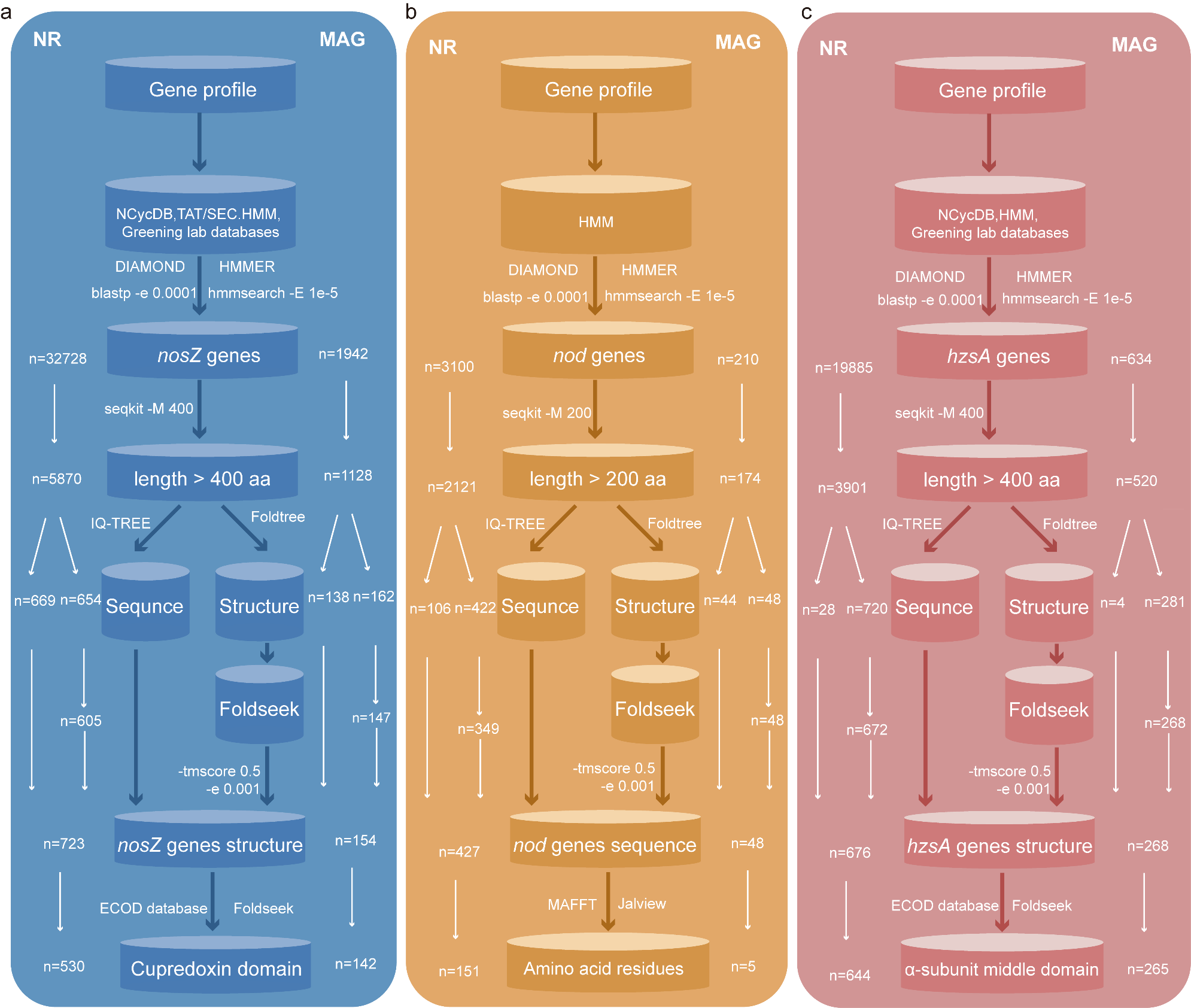

**Supplementary** **Figure 4. Workflow for** **the identification and validation of *nosZ* (a), *nod* (b) and *hzsA* (c) sequences.** The workflow is initiated with metagenomic data derived from 165 cold seep samples. This data is then bifurcated into two processing streams: one involving 373 million protein sequences from metagenomic contigs condensed to 147 million non-redundant genes (NR), and the other involving 6.29 million genes predicted from 3,164 dereplicated species-level MAGs via Prodigal. The non-redundant genes and predicted genes from MAGs are further examined against different databases, and filtered using stringent selection criteria including sequence length, identity, TM-score. This selection process yields a set of candidates NosZ, NOD and HzsA sequences which are subjected to phylogenetic and structural tree, and the presence of motifs or protein active domains for final confirmation, resulting in validated NosZ (NR: 530; MAG: 142), NOD (NR: 151; MAG: 5) and HzsA (NR: 644; MAG: 265) sequences respectively.

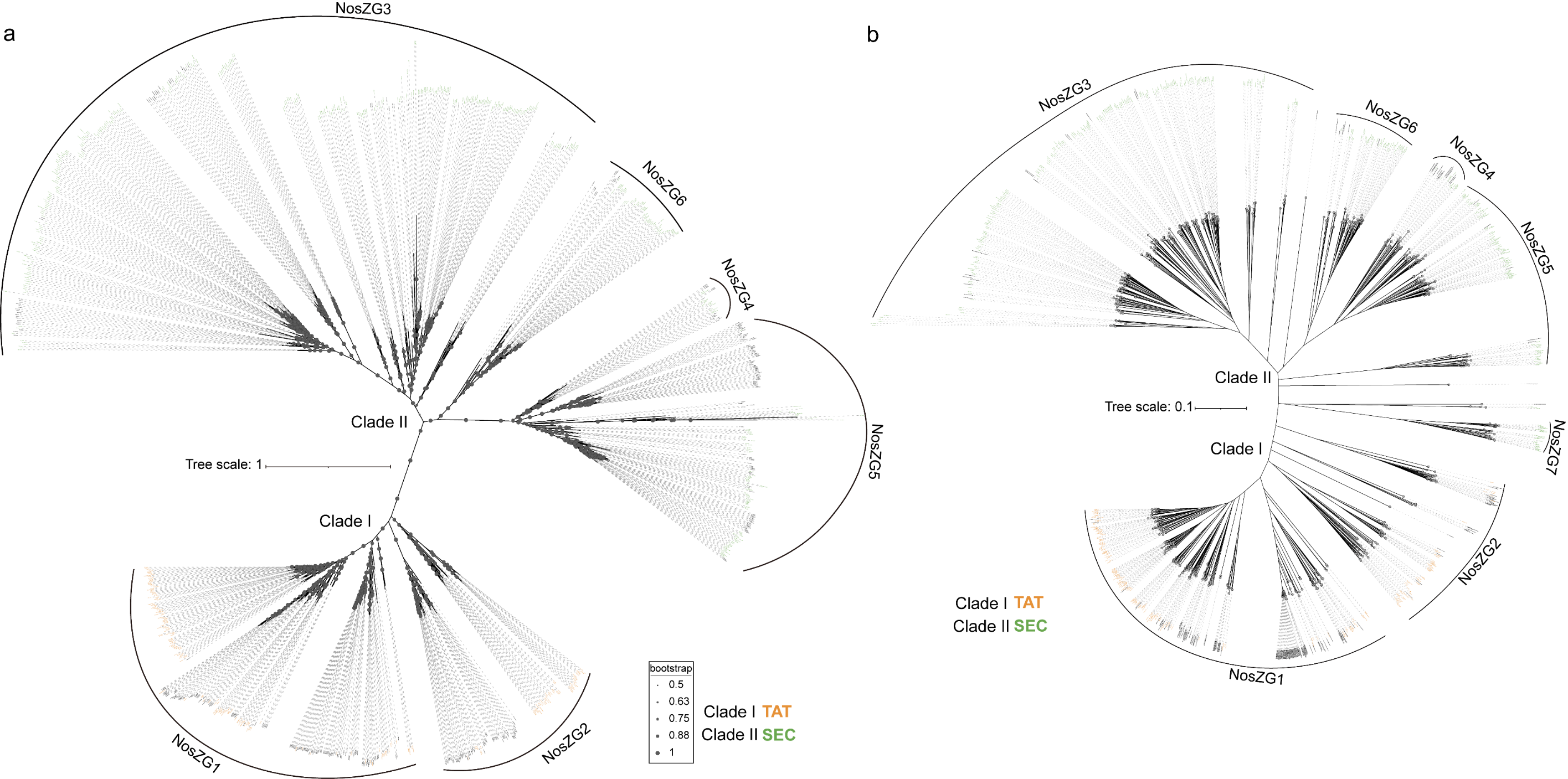

**Supplementary Figure 5**. **Maximum-likelihood phylogenetic trees of** ***nosZ* protein sequences (a) and structure trees of *nosZ* protein sequences with Foldtree (b) recovered from 147 million non-redundant genes (NR)**. Each group of *nosZ* is labeled by a unique color: *nosZ* Clade I, orange; *nosZ* Clade II, green. Scale bar indicates the mean number of substitutions per site. Bootstrap values over 50% were shown next to the nodes in phylogenetic trees. A sequence-divergent branch NosZG7 (n = 27) from *nosZ* Clade II was only found through structural analysis.

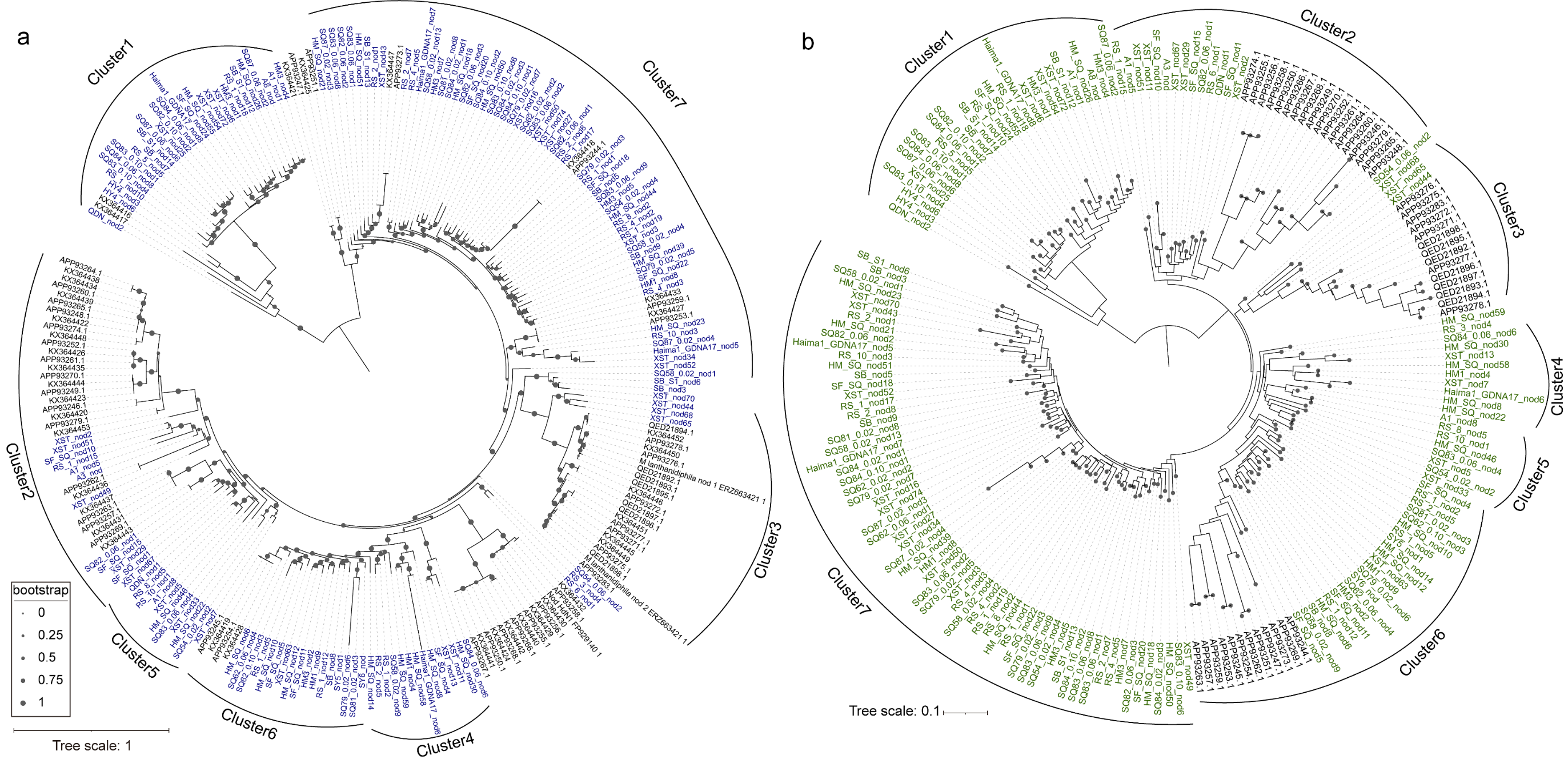

**Supplementary Figure 6**. **Maximum-likelihood** **phylogenetic trees of *nod* protein sequences (a) and structure trees of *nod* protein sequences with Foldtree (b) recovered from 147 million non-redundant genes (NR).** Each *nod* is labeled by dark blue and dark green in phylogenetic trees and structure trees, respectively. Scale bar indicates the mean number of substitutions per site. Bootstrap values over 50% were shown next to the nodes in phylogenetic trees. The sequence-divergent branches Cluster1 (n = 28) and Cluster5 (n = 8) were only found through phylogenetic and structural analysis, respectively.

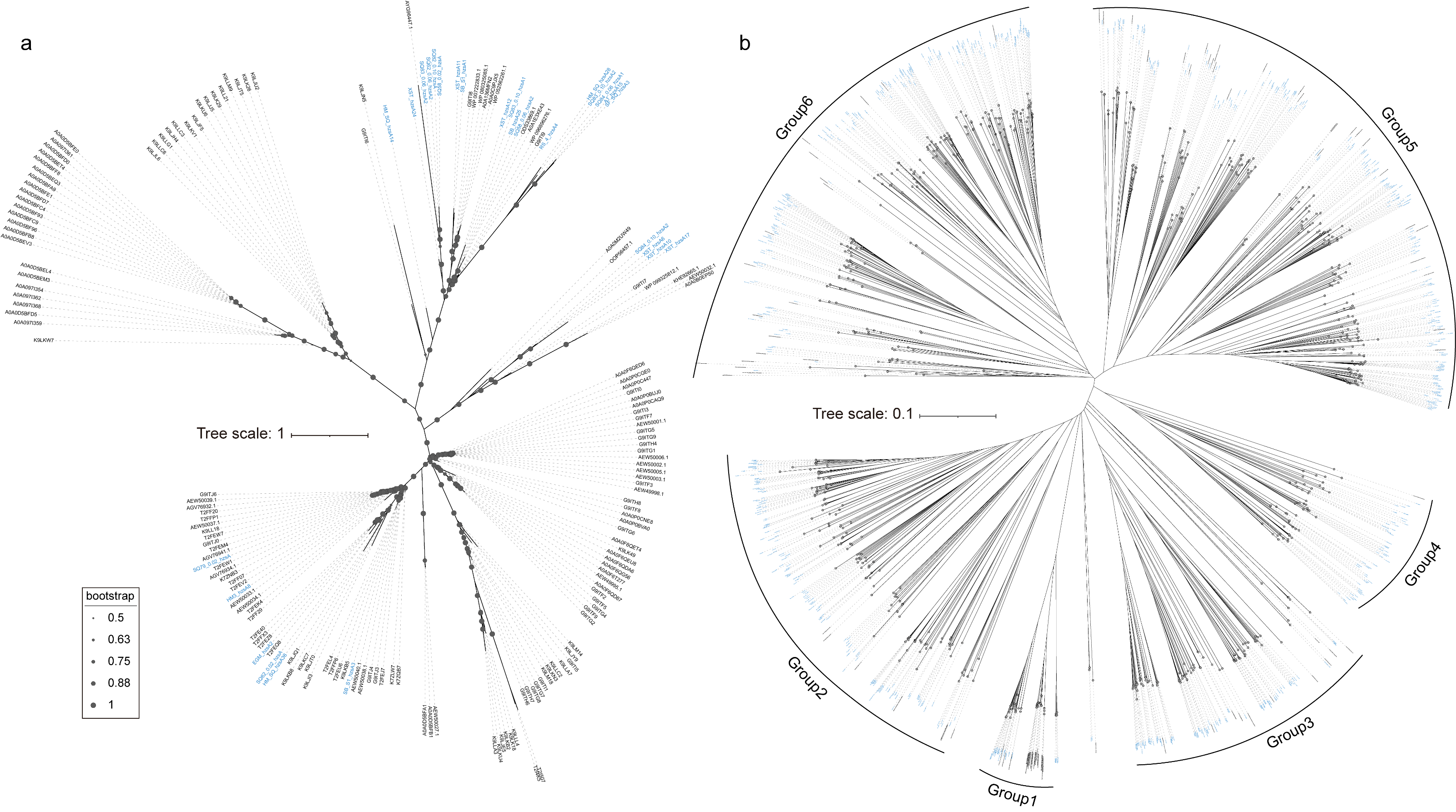

**Supplementary Figure 7**. **Maximum-likelihood phylogenetic trees of *hzsA* protein sequences (a) and structure trees of *hzsA* protein sequences with Foldtree (b) recovered from 147 million non-redundant genes (NR).** Each cold seep-derived *hzsA* sequence is labeled by blue in the phylogenetic tree and structure tree. Scale bars indicate the mean number of substitutions per site. Bootstrap values over 50% were shown next to the nodes in phylogenetic trees.

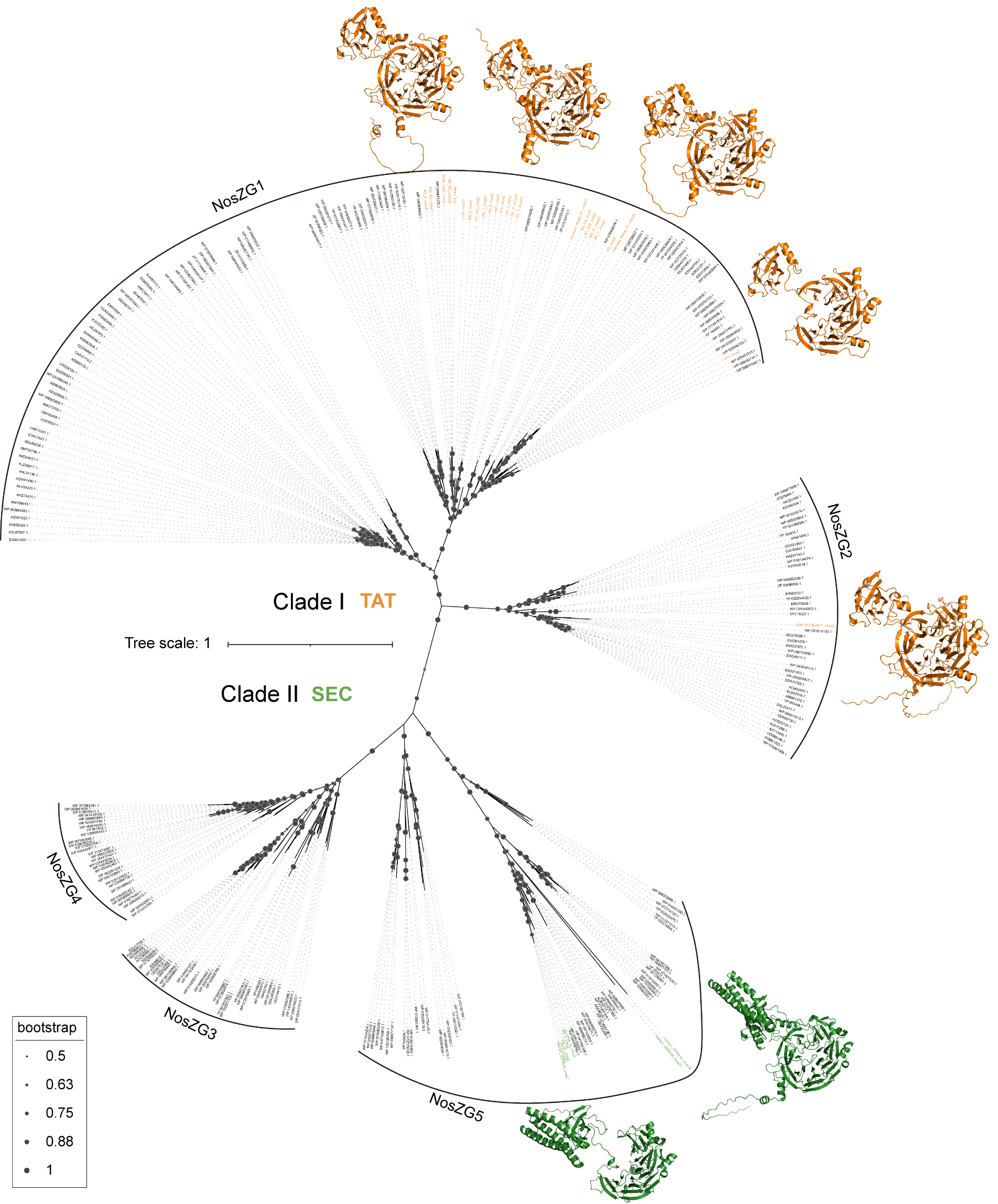

**Supplementary Figure 8. Maximum-likelihood phylogenetic trees of *nosZ* protein sequences in plasmid and structure of *nosZ* protein in plasmid with Alphafold2.** Each group of *nosZ* is labeled by a unique color: *nosZ* Clade I, orange; *nosZ* Clade II, green. Scale bar indicates the mean number of substitutions per site. Bootstrap values over 50% were shown next to the nodes in phylogenetic trees. Surrounding the tree, the AlphaFold-predicted 3D structures of representative NosZ are displayed.

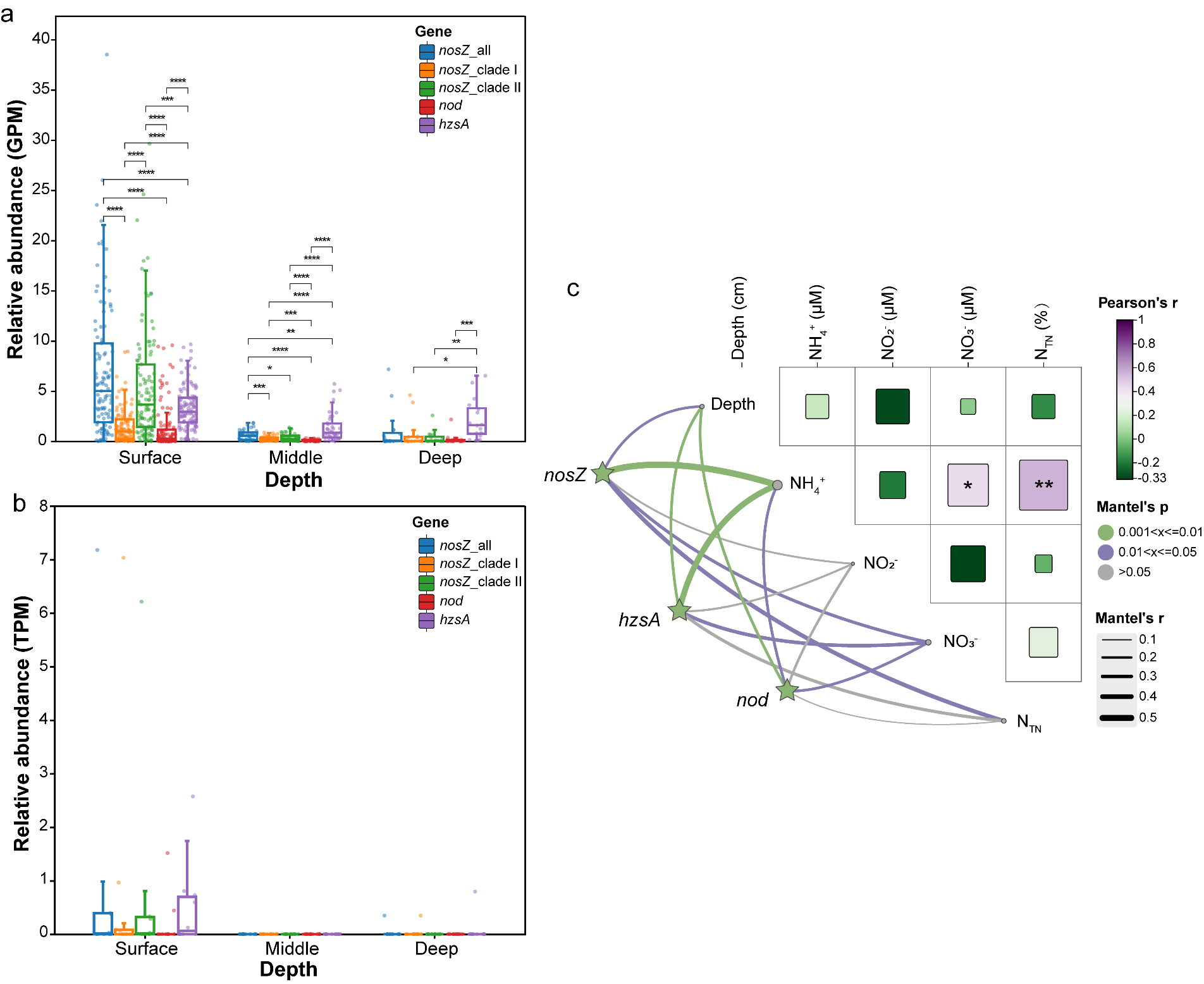

**Supplementary Figure 9. The relative abundance and expression levels of nitrogen-loss genes at different depth of cold seeps.** (a) Relative abundance measured as GPM for metagenomes. (b) Expression levels measured as TPM for transcripts. Inserted plots show the all N-loss genes abundance: all *nosZ* genes are shown in dark blue, *nosZ* Clade I genes in orange, *nosZ* Clade II genes in green, *nod* genes in dark red, *hzsA* genes in dark purple. *P* values of differences across different types of N-loss genes were computed through Kruskal-Wallis rank-sum tests. (c) Pairwise comparisons of environmental factors are shown, with a color gradient denoting Pearson’s correlation coefficient. Relative abundance (GPM) of *nosZ*, *nod* and *hzsA* genes was related to each environmental factor by Mantel tests. Edge width corresponds to the Mantel’s r statistic, and edge color denotes the statistical significance based on 9,999 permutations. Detailed data and statistics for significance tests are provided in **Supplementary Tables 4-7**.

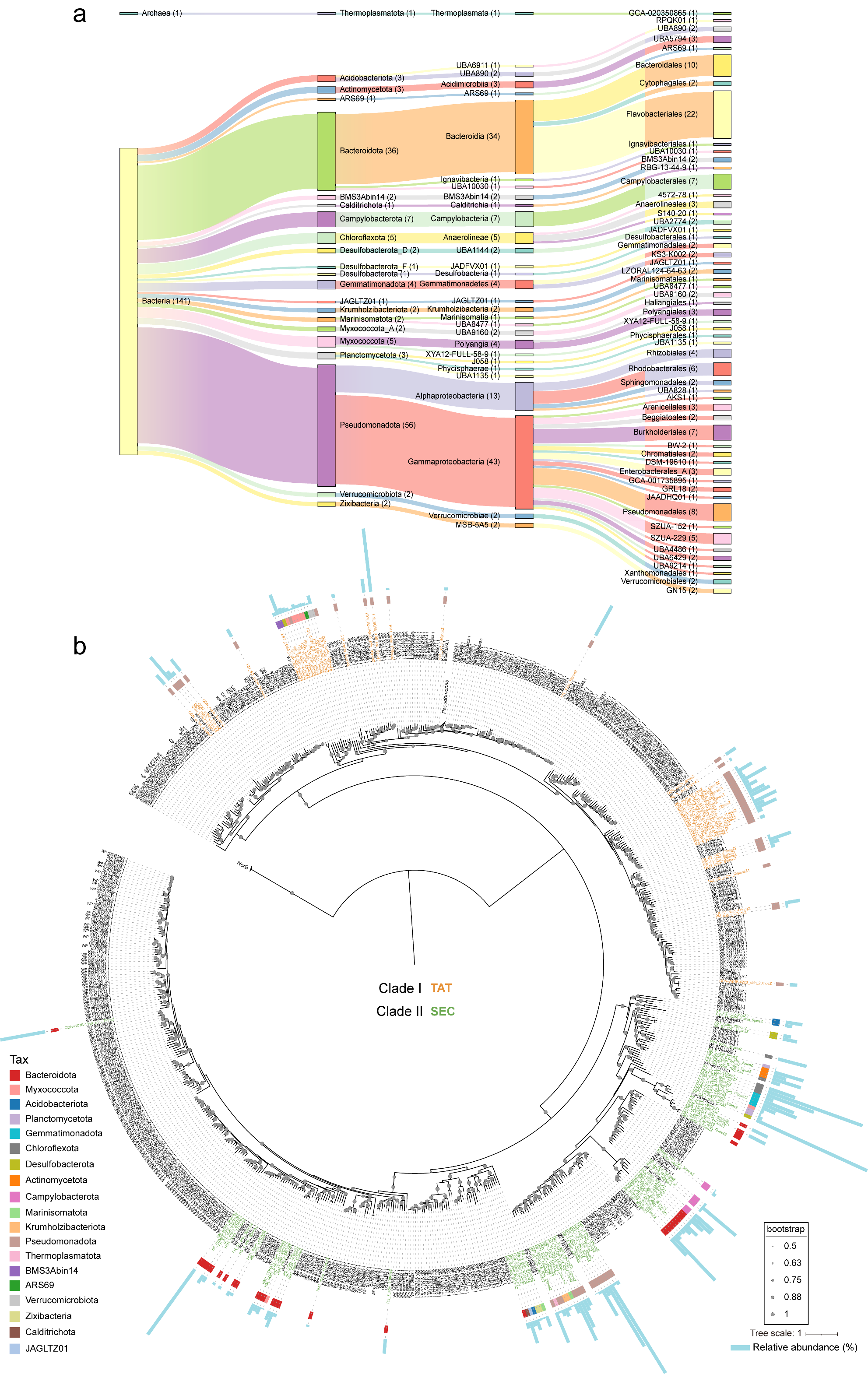

**Supplementary Figure 10**. **Taxonomic classification and maximum-likelihood phylogenetic trees of *nosZ* protein sequences** **recovered from 3,164 cold seep MAGs.** (a) A Sankey diagram showing the taxonomy of archaeal and bacterial MAGs according to GTDB, categorizing nitrous-oxide reducers across taxonomic levels. (b) Maximum-likelihood phylogenetic trees of *nosZ* protein sequences. Each group of *nosZ* is labeled by a unique color: *nosZ* Clade I, orange; *nosZ* Clade II, green. Scale bar indicates the mean number of substitutions per site. Bootstrap values over 50% were shown next to the nodes in phylogenetic trees. Surrounding the tree, the microbial compositions and relative abundance of *nosZ* in MAGs are displayed. Details for microbial compositions and relative abundance of *nosZ* are provided in **Supplementary Table 8**.

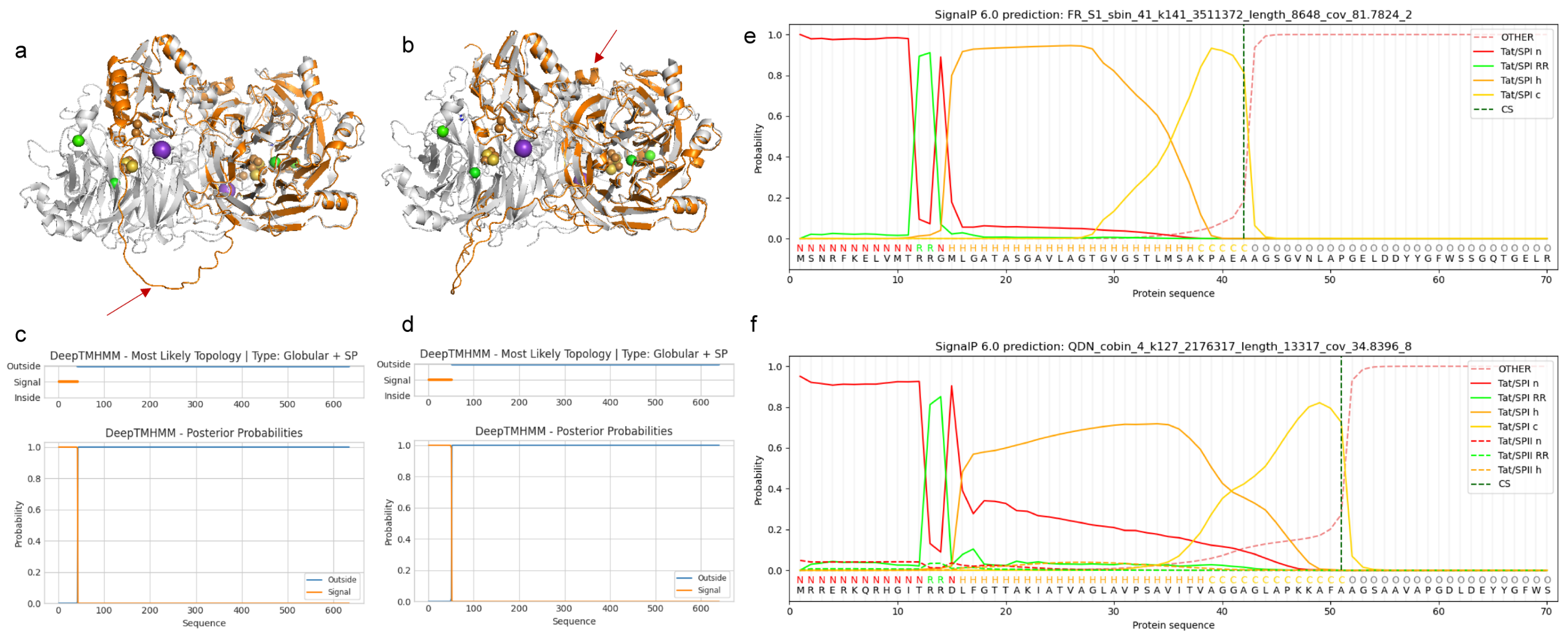

**Supplementary** Figure 11**. Structural alignments and topological structure prediction of** **NosZs in clade I.** Structural alignments of the NosZ of FR_S1_sbin_41_k141_3511372_length_8648_cov_81.7824_2 gene (a) and QDN_cobin_4_k127_2176317_length_13317_cov_34.8396_8 gene (b) belonging to *Alphaproteobacteria* (*Pseudomonadota*). The enzyme is a homodimer in head-to-tail orientation with the tetranuclear Cu_Z_ active site located in the N-terminal, seven-bladed b-propeller domain and the binuclear Cu_A_ site in the C-terminal cupredoxin domain. The NosZs in clade I recovered from 3,164 cold seep MAGs are shown in orange, and reference NosZ in gray from *Pseudomonas stutzeri* nitrous oxide reductase with 3SBQ in RCSB Protein Data Bank (RCSB PDB). The enzymes greatly vary in their N-terminal and C-terminal structures highlighted with a red arrow. Topological structure prediction with DeepTMHMM of the NosZ of FR_S1_sbin_41_k141_3511372_length_8648_cov_81.7824_2 gene (c) and QDN_cobin_4_k127_2176317_length_13317_cov_34.8396_8 gene (d). Topological structure prediction with SignalP of the NosZ of FR_S1_sbin_41_k141_3511372_length_8648_cov_81.7824_2 gene (e) and QDN_cobin_4_k127_2176317_length_13317_cov_34.8396_8 gene (f). Detailed data for topological structure prediction with SignalP of *nosZ* are provided in **Supplementary Table 14**.

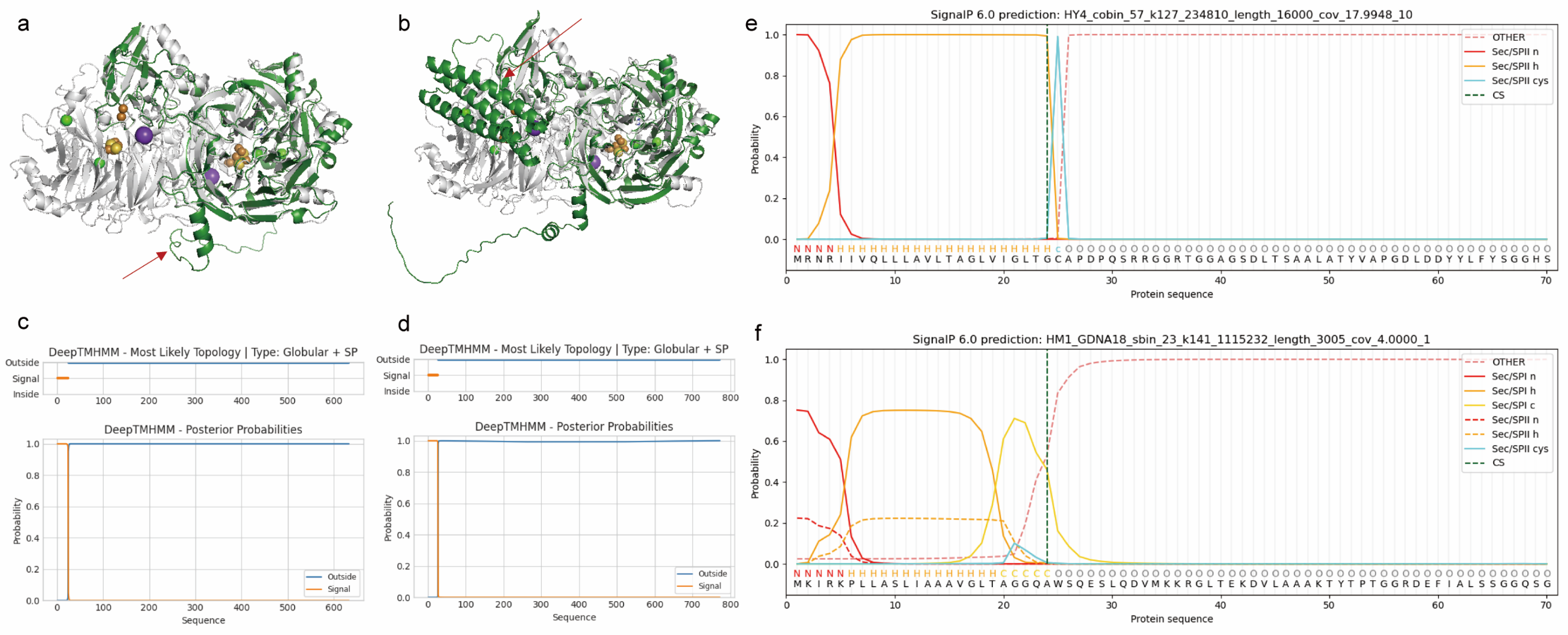

**Supplementary** Figure 12**. Structural alignments and topological structure prediction of NosZs in clade II.** Structural alignments of the NosZ of HY4_cobin_57_k127_234810_length_16000_cov_17.9948_10 gene (a) and HM1_GDNA18_sbin_23_k141_1115232_length_ 3005_cov_4.0000_1 gene (b) belonging to UBA6911 (*Acidobacteriota*) and *Thermoplasmata* (*Thermoplasmatota*). The enzyme is a homodimer in head-to-tail orientation with the tetranuclear Cu_Z_ active site located in the N-terminal, seven-bladed b-propeller domain and the binuclear Cu_A_ site in the C-terminal cupredoxin domain. The NosZs in clade II recovered from 3,164 cold seep MAGs are shown in orange, and reference NosZ in gray from *Pseudomonas stutzeri* nitrous oxide reductase with 3SBQ in RCSB Protein Data Bank (RCSB PDB). The enzymes greatly vary in their N-terminal and C-terminal structures highlighted with a red arrow. Topological structure prediction with DeepTMHMM of the NosZ of HY4_cobin_57_k127_234810_length_16000_cov_17.9948_10 gene (c) and HM1_GDNA18_sbin_23_k141_1115232_length_3005 _cov_4.0000_1 gene (d). Topological structure prediction with SignalP of the NosZ of HY4_cobin_57_k127_234810_length_16000_cov _17.9948_10 gene (e) and HM1_GDNA18_sbin_23_k141_1115232_length_3005_cov_4.0000_1 gene (f). Detailed data for topological structure prediction with SignalP of *nosZ* are provided in **Supplementary Table 14**.

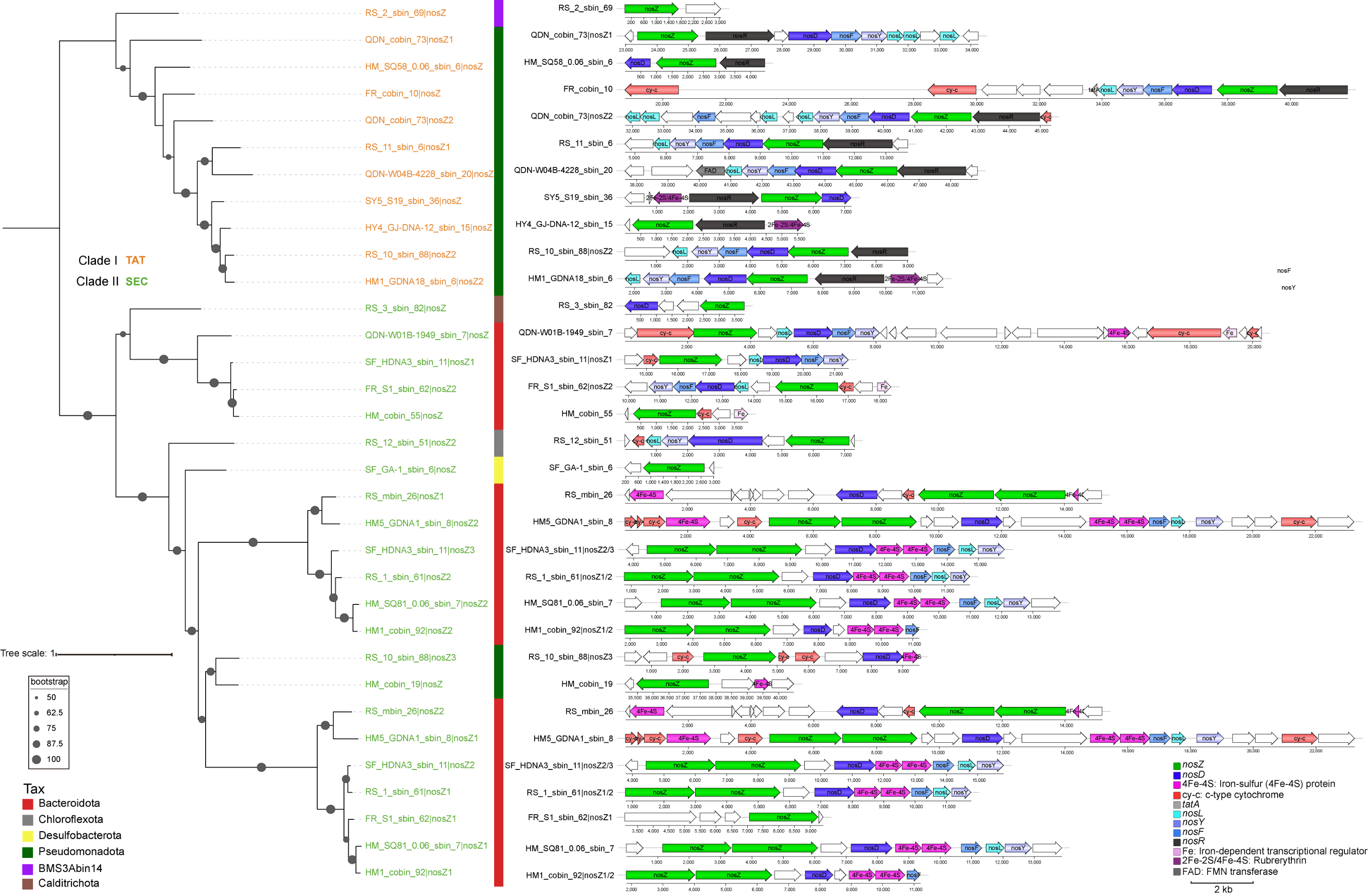

**Supplementary** Figure 13**.** **Comparison of *nos* clusters from genomes harboring Clade I or Clade II *nosZ* genes.** Phylogeny at left is based on the phylogeny shown in Figure 5 with phylum level taxonomic designation, and the scale bar at right indicating estimated amino-acid substitutions per site. *nos* clusters of Clade II harbor the atypical *nosZ* and encode predicted iron-sulfur-binding proteins (labeled “4Fe-4S” or “2Fe-2S”), *c*-type cytochromes (cy-c), or ABC transporter proteins (labeled “ABC” or “ABC-2”). Accessory genes (*nosD*, *nosF*, *nosL*, and *nosY*) are generally conserved across *no*s clusters with both typical (Clade I) and atypical *nosZ*. Noncolored genes in the operons have no orthologs in any other known *nos* cluster. *nosR* and *nosX* are associated exclusively with typical *nos* clusters. Details for *nosZ*-containing contig annotations are provided in **Supplementary Tables 12-13**.

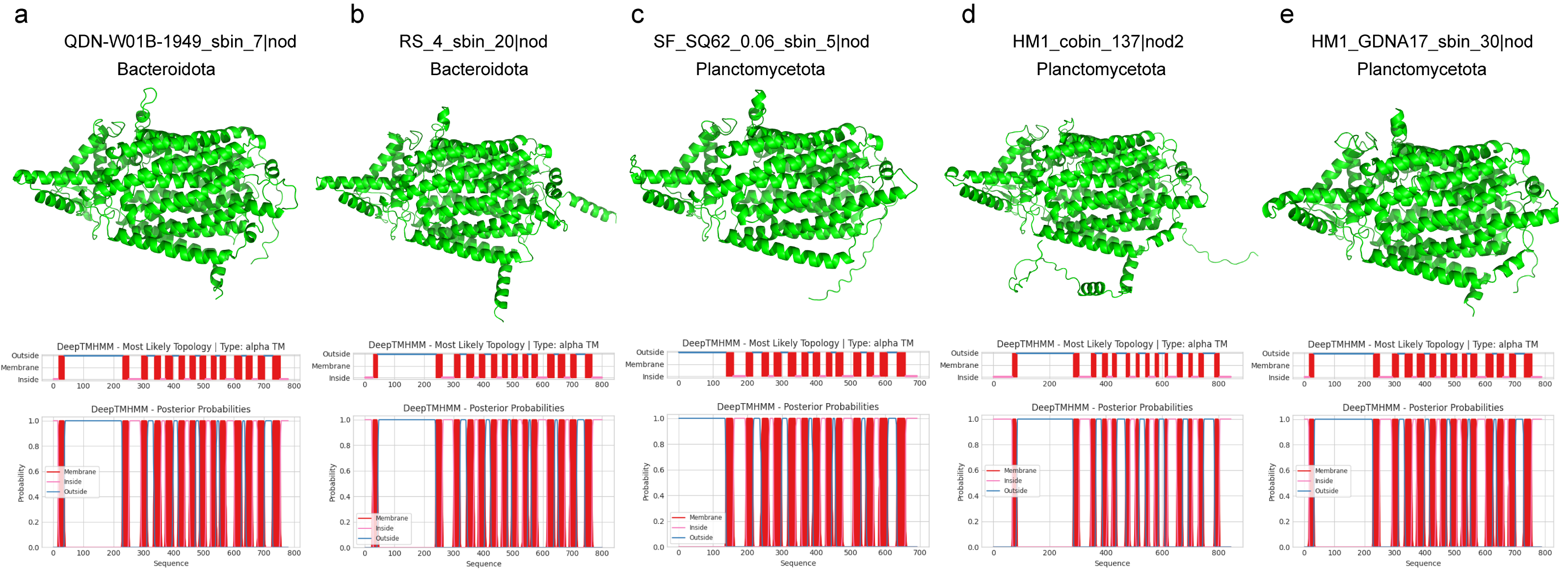

**Supplementary** Figure 14**. Structure of *nod* protein recovered from 3,164 cold seep MAGs.** (a, b) Structure of *nod* protein in MAGs belonging to *Bacteroidota* with AlphaFold2 and topological structure prediction with DeepTMHMM. (c-e) Structure of *nod* protein in MAGs belonging to *Planctomycetota* with AlphaFold2 and topological structure prediction with DeepTMHMM. Detailed data for topological structure prediction with DeepTMHMM of *nod* are provided in **Supplementary Table 16**.

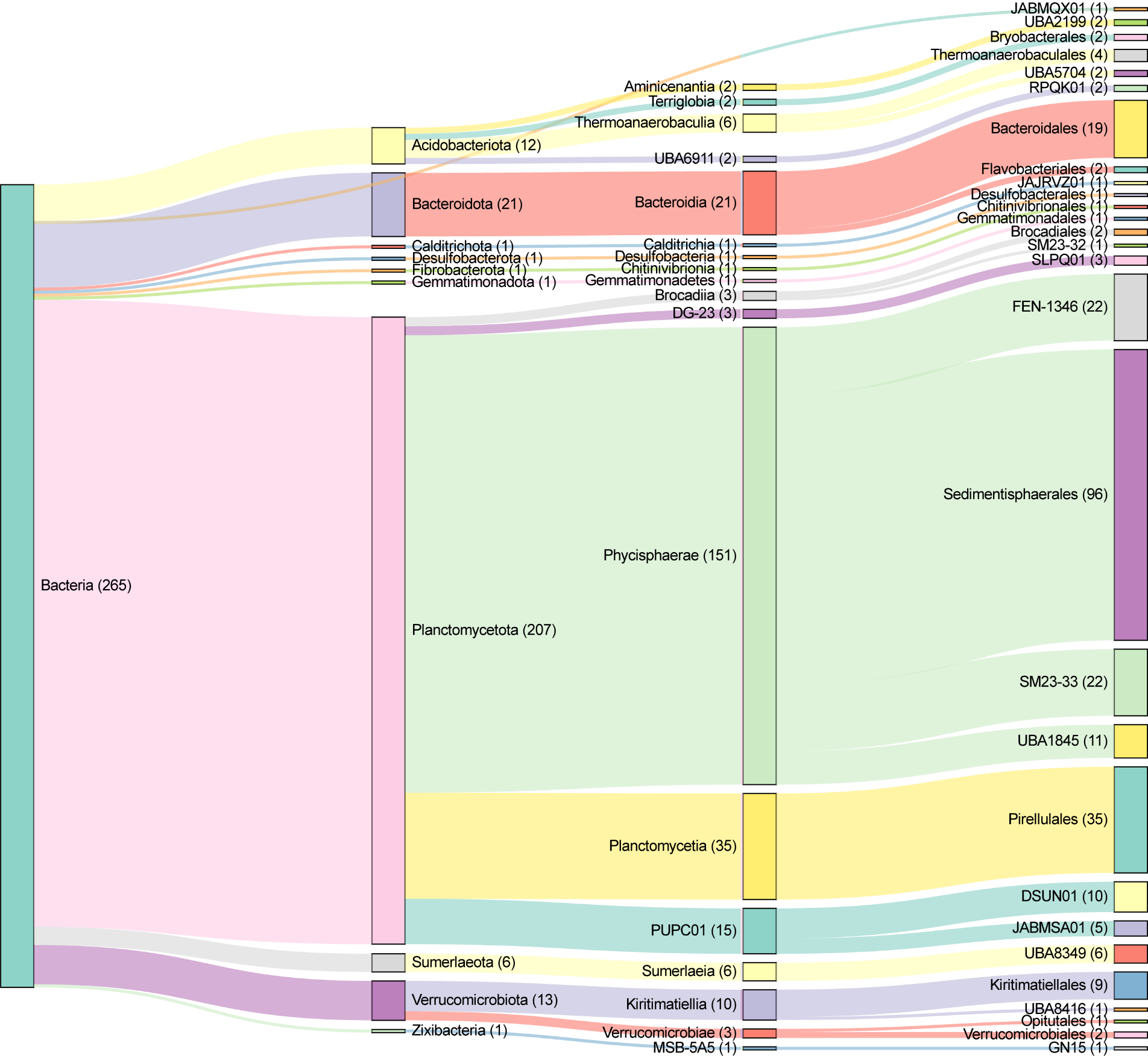

**Supplementary Figure 15**. **Taxonomic classification of *hzsA* protein sequences recovered from 3,164 cold seep MAGs.** A Sankey diagram showing the taxonomy of MAGs according to GTDB, categorizing anammox bacterium across taxonomic levels. Detailed for taxonomic classification are provided in **Supplementary Table 18**.

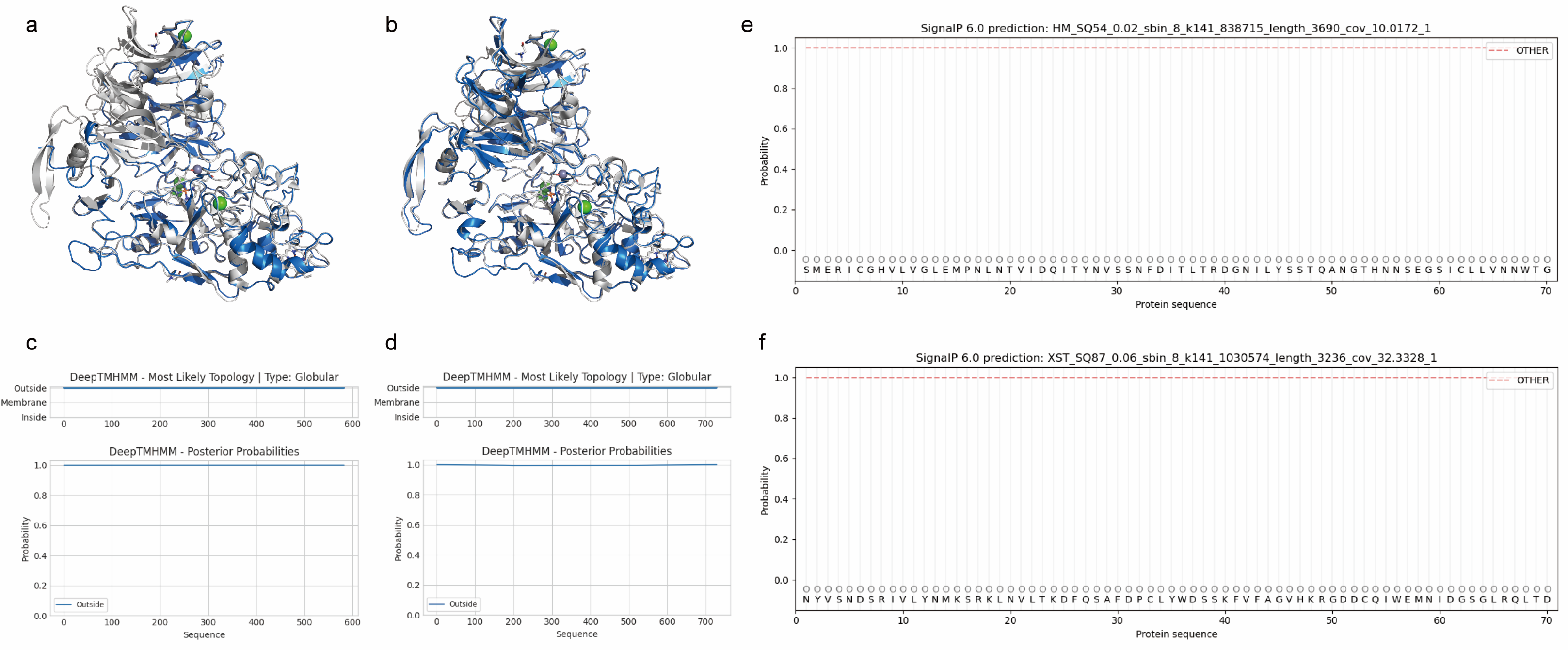

**Supplementary** Figure 16**. Structural alignments and topological structure prediction of traditional *hzsA* protein in MAGs.** Structural alignments of the HzsA of HM_SQ54_0.02_sbin_8_k141_838715_length_3690_cov_10.0172_1 gene (a) and XST_SQ87_0.06_sbin _8_k141_1030574_length_3236_cov_32.3328_1 gene (b) belonging to *Scalinduaceae* (*Planctomycetota*). The the HzsA with two haem groups contains N-terminal domain, middle domain and C-terminal domain. The traditional HzsA recovered from 3,164 cold seep MAGs are shown in dark blue, and reference HzsA in gray from *Candidatus Kuenenia stuttgartiensis* hydrazine synthase with 5C2V in RCSB Protein Data Bank (RCSB PDB). Topological structure prediction with DeepTMHMM of the HzsA of HM_SQ54_0.02_sbin_8_k141_838715_length_3690_cov _10.0172_1 gene (c) and XST_SQ87_0.06_sbin_8_k141_1030574_length_3236_cov _32.3328_1 gene (d). Topological structure prediction with SignalP of the NosZ of HM_SQ54_0.02_sbin_8_k141_838715_length_3690_cov_10.0172_1 gene (e) and XST_SQ87_0.06 _sbin_8_k141_1030574_length_3236_cov_32.3328_1 gene (f).

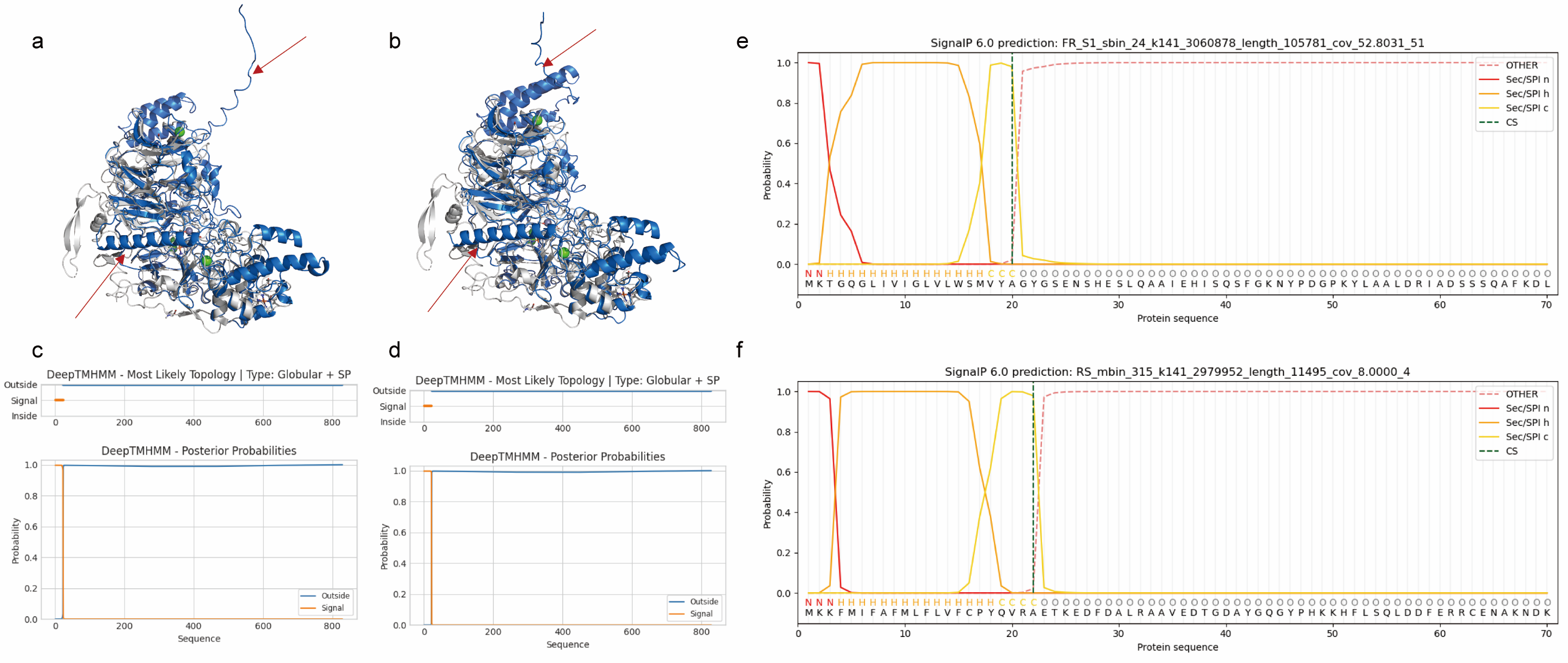

**Supplementary** Figure 17**. Structural alignments and topological structure prediction of novel *hzsA* protein in MAGs.** Structural alignments of the HzsA of FR_S1_sbin_24_k141_3060878_length_105781_cov_52.8031_51 gene belonging to *Verrucomicrobiales* (*Verrucomicrobiota*) (a) and RS_mbin_315_k141_2979952_length_11495_cov_8.0000_4 gene belonging to *Anaerohalosphaera* (*Planctomycetota*) (b). The the HzsA with two haem groups contains N-terminal domain, middle domain and C-terminal domain. The traditional HzsA recovered from 3,164 cold seep MAGs are shown in dark blue, and reference HzsA in gray from *Candidatus Kuenenia stuttgartiensis* hydrazine synthase with 5C2V in RCSB Protein Data Bank (RCSB PDB). The C-terminal structures and N-terminal transmembrane *α*-helix contained in novel HzsA highlighted with a red arrow. Topological structure prediction with DeepTMHMM of the HzsA of FR_S1_sbin_24_k141_3060878_length_105781_cov_52.8031_51 gene (c) and RS_mbin_315_k141_2979952_length_11495_cov_8.0000_4 gene (d). Topological structure prediction with SignalP of the NosZ of FR_S1_sbin_24_k141_3060878_length_105781_cov_52.8031_51 gene (e) and RS_mbin_315_k141_2979952_length_11495_cov_8.0000_4 gene (f).

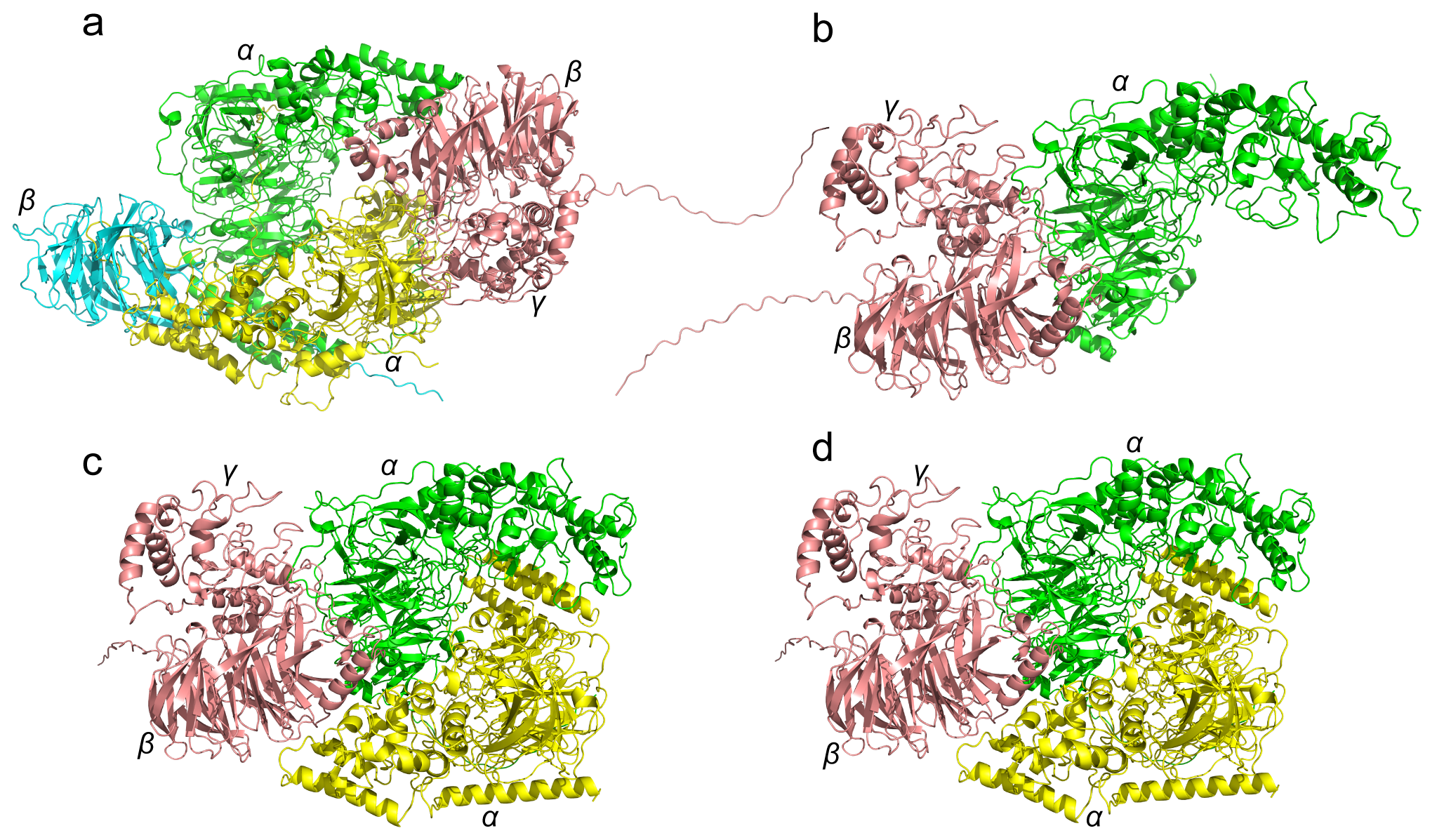

**Supplementary** Figure 18**. Predicted structure of HZS in representative MAGs.** Predicted structure of HZS in QDN-W03B-2880_sbin_17 belonging to FEN-1346 (*Planctomycetota*) (a), RS_18_sbin_51 belonging to *Sedimentisphaerales* (*Planctomycetota*) (b), RS_mbin_315 belonging to *Sedimentisphaerales* (*Planctomycetota*) (c), and QDN-W04B-4228_sbin_32 belonging to *Bacteroidales* (*Bacteroidota*) (d). HZS complex structure; *α*-subunits are coloured green and yellow, *β*-subunits are blue and brick red, and *γ*-subunits are grey and rose red.
